# Polo-like kinase phosphorylation of the orphan kinesin KIN-G negatively regulates centrin arm biogenesis in *Trypanosoma brucei*

**DOI:** 10.64898/2026.01.16.699953

**Authors:** Yasuhiro Kurasawa, Qing Zhou, Kyu Joon Lee, Huiqing Hu, Ziyin Li

## Abstract

The unicellular parasite *Trypanosoma brucei* assembles a motile flagellum that is required for locomotion, cell division plane placement, and cell-cell communication. Inheritance of the flagellum during the cell cycle relies on the faithful duplication/segregation of multiple flagellum-associated cytoskeletal structures, including a centrin-marked, bar-shaped structure termed centrin arm, which also determines the site for Golgi assembly. Biogenesis of the centrin arm requires the Polo-like kinase homolog TbPLK and the orphan kinesin KIN-G, but the mechanistic role of TbPLK in centrin arm biogenesis remains elusive. Here we report that TbPLK phosphorylates KIN-G, disrupts its microtubule-binding activity, and negatively regulates its function. TbPLK phosphorylates KIN-G *in vitro* at multiple residues, two of which are *in vivo* TbPLK phosphosites, including the Thr301 residue within one of the microtubule-binding motifs of the kinesin motor domain. Phosphorylation of Thr301 by TbPLK inhibits the microtubule-binding activity of KIN-G *in vitro*, and expression of a Thr301 phospho-mimic mutant in *T. brucei* disrupts centrin arm integrity, Golgi duplication, flagellum attachment zone elongation, flagellum positioning, and cell division plane placement. In wild-type *T. brucei* cells, Thr301 phosphorylation occurs on a small portion of the KIN-G population, suggesting that KIN-G undergoes phosphorylation/dephosphorylation cycles to regulate its activity. Together, these findings uncover a negative role of TbPLK-mediated phosphorylation of KIN-G in regulating centrin arm biogenesis in trypanosomes.

## Introduction

The early branching parasitic protozoan *Trypanosoma brucei* causes sleeping sickness in humans and nagana in cattle in sub-Saharan Africa. This parasite possesses a motile flagellum, which is required for cell motility, cell division, and cell-cell communication. The flagellum originates from the basal body, which has a typical structure of centrioles in eukaryotes, exits the cell body through the flagellar pocket, and extends toward the anterior tip of the cell by attaching to the cell membrane via the flagellum attachment zone (FAZ), a specialized cytoskeletal structure consisting of an intra-cellular FAZ filament, a flagellum FAZ domain, and an inter-membrane junction between the flagellum and the cell body (Sunter and Gull, 2016). At the proximal end of the flagellum near the flagellar pocket region, there sit several flagellum-associated cytoskeletal structures, including the flagellar pocket collar (FPC) (Lacomble et al., 2009) and the hook complex (Esson et al., 2012). The horseshoe-shaped FPC, which is marked by TbBILBO1, wraps around the flagellum, whereas the hairpin-shaped hook complex, which consists of a fishhook-like structure marked by TbMORN1 and a bar-shaped centrin arm structure marked by TbCentrin2 (Esson et al., 2012; Morriswood, 2015), sits on the top of the FPC (Esson et al., 2012). Between the mature basal body and the pro-basal body, there originates a specialized set of four microtubules termed the microtubule quartet (MtQ), which passes through the hook complex and the FPC and extends, alongside the intracellular FAZ filament, to the anterior tip of the cell (Esson et al., 2012; Vaughan and Gull, 2016). The intracellular FAZ filament originates from a region within the hook complex structure (Esson et al., 2012) and elongates alongside the MtQ toward the anterior cell tip (Sunter et al., 2015; Zhou et al., 2015). In close proximity to the centrin arm, the Golgi apparatus is located (He et al., 2004), and it sits next to the endoplasmic reticulum exit site (ERES), forming unique junctions to facilitate protein trafficking (Bangs, 2011).

During the S phase of the cell cycle in *T. brucei*, a new pair of mature basal body/pro-basal body is assembled, followed by the assembly of a new flagellum from the mature basal body and the subsequent assembly of the flagellum-associated cytoskeletal structures. The newly synthesized flagellum and its associated, newly synthesized cytoskeletal structures are segregated into the new-flagellum daughter (NFD) cell, whereas the old-flagellum daughter (OFD) cell inherits the existing flagellum and its associated cytoskeletal structures (Abeywickrema et al., 2019). Segregation or positioning of the newly assembled flagellum depends on the faithful duplication and segregation of its associated cytoskeletal structures, which further impacts the placement of the cell division plane/cleavage furrow and the faithful division of the cell (Hu et al., 2015; Pham et al., 2020; Zhou et al., 2010; Zhou et al., 2011). During the cell cycle, the Golgi apparatus undergoes *de novo* duplication, assembling a new Golgi next to the old Golgi in a centrin arm-dependent manner (He et al., 2005), and the newly assembled Golgi is segregated into the NFD cell, likely by associating with the newly assembled centrin arm through the scaffold protein CAAP1 (Zhou et al., 2024).

The precise function of the hook complex remains elusive, but its deficiency often leads to defects in FAZ elongation, flagellum positioning/segregation, and cell division plane placement (Pham et al., 2020; Pham et al., 2019; Zhou et al., 2010; Zhou et al., 2025; Zhou et al., 2024). Several regulatory proteins that control hook complex biogenesis have been identified, including the Polo-like kinase homolog TbPLK (de Graffenried et al., 2008), the kinetoplastid-specific protein phosphatase KPP1 (Zhou et al., 2018b), the E3 ubiquitin ligase complex CRL4^WDR1^ (Hu et al., 2017), the orphan kinesin KIN-G (Zhou et al., 2024), and the KIN-G-interacting protein WDR2 (Zhou et al., 2025). TbPLK phosphorylates the centrin arm protein TbCentrin2 to promote centrin arm duplication (de Graffenried et al., 2013), whereas its protein abundance is regulated by CRL4^WDR1^, which also promotes centrin arm biogenesis (Hu et al., 2017). KIN-G is a microtubule plus end-directed motor protein, which localizes to centrin arm and regulates centrin arm biogenesis (Zhou et al., 2024), and WDR2 promotes centrin arm biogenesis by recruiting KIN-G (Zhou et al., 2025). KIN-G is phosphorylated at multiple residues *in vivo* in trypanosome cells (Nett et al., 2009; Zhou et al., 2025), but the protein kinase responsible for KIN-G phosphorylation was not identified previously and whether phosphorylation of KIN-G may regulate KIN-G motor activity and/or cellular function remains unknown.

Here we report that TbPLK phosphorylates KIN-G *in vitro* and *in vivo* at two sites, the Thr301 residue in one of the microtubule-binding motifs of the motor domain and the Ser569 residue within the third coiled-coil motif in the C-terminus. Phosphorylation of Thr301 inhibits the microtubule-binding activity of KIN-G, and expression of the Thr301 phospho-mimic mutant in trypanosomes disrupts KIN-G’s cellular function. These results suggest that TbPLK negatively regulates KIN-G activity through phosphorylating Thr301 and that dephosphorylation of Thr301 is required for KIN-G to exert its cellular function. This finding underscores a novel control mechanism of KIN-G by Polo-like kinase in regulating centrin arm biogenesis in *T. brucei*.

## Results

### TbPLK co-localizes with KIN-G at centrin arm and phosphorylates KIN-G *in vitro* and *in vivo*

The findings that TbPLK and KIN-G are both required for the biogenesis of the centrin arm and its associated Golgi apparatus (de Graffenried et al., 2008; Zhou et al., 2024) and that KIN-G is phosphorylated *in vivo* in trypanosome cells (Nett et al., 2009; Zhou et al., 2025) prompted us to test whether TbPLK may phosphorylate KIN-G and thus may regulate KIN-G function. We first investigated when the two proteins may co-localize during the cell cycle by co-immunofluorescence microscopy analysis of cells expressing endogenously triple HA-tagged KIN-G. During G1 phase, TbPLK and KIN-G co-localized at the centrin arm, with KIN-G more enriched at the distal end of the centrin arm (Figure 1A). During early S-phase when the centrin arm started to duplicate, KIN-G and TbPLK co-localized, almost entirely, to the elongating centrin arm (Figure 1A, S-phase cell on the left). When cells progressed to late S-phase, the centrin arm was duplicated and separated to two adjacent structures, KIN-G localized to the separated centrin arm structures (Figure 1A, S-phase cell on the right). However, during this stage, TbPLK was enriched at the new FAZ tip and the flagella connector, with a small amount of TbPLK remaining on the old centrin arm, where it co-localized with KIN-G (Figure 1A, S-phase cell on the right). During G2 phase when the two centrin arms were farther separated, KIN-G remained on the two centrin arm structures, but TbPLK no longer resided on the centrin arm, but was enriched on the new FAZ tip and the flagella connector (Figure 1A). Note that from G1 phase to G2 phase, TbPLK was additionally localized to the basal body (Figure 1A).

**Figure 1.**
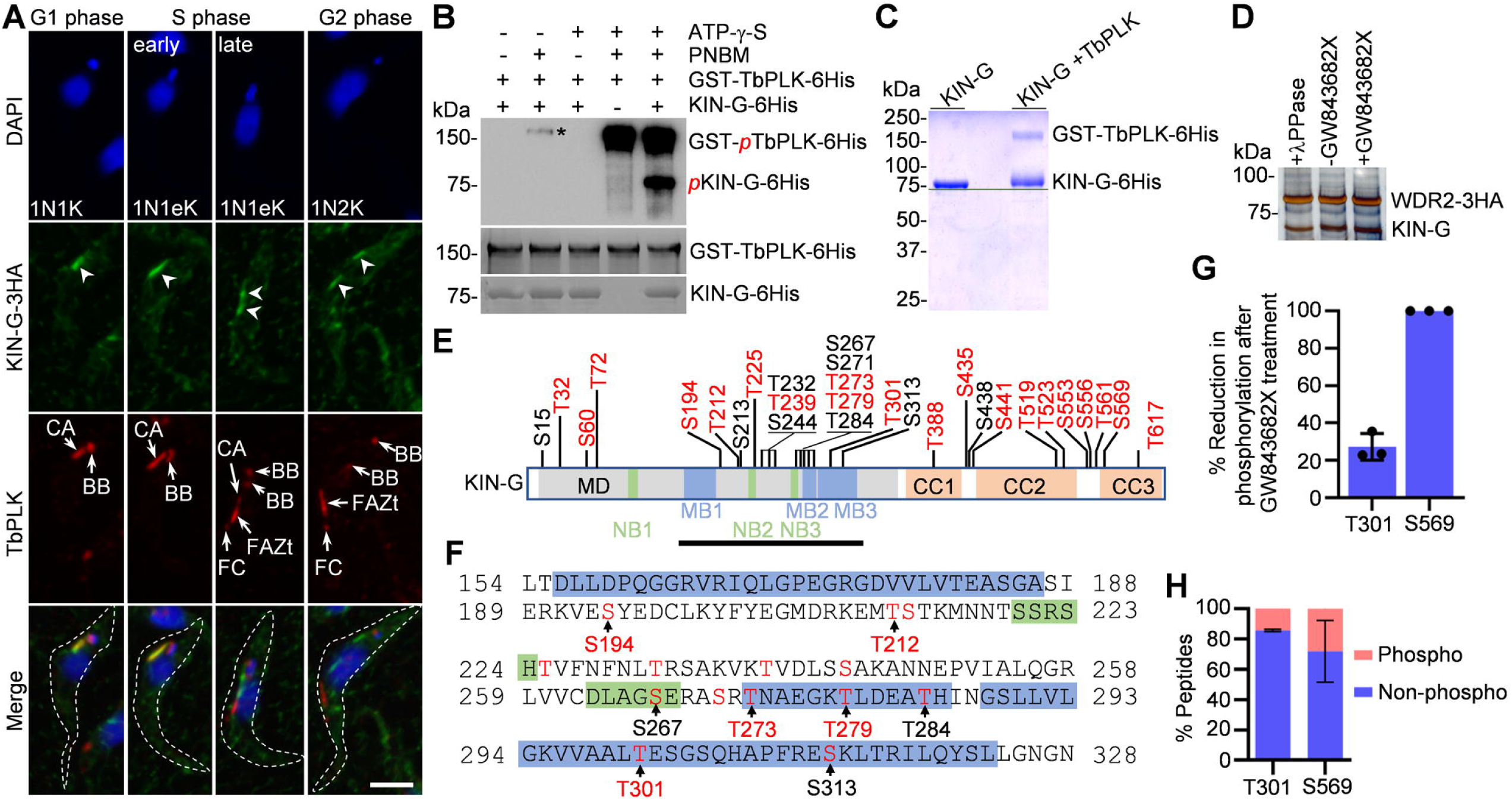
KIN-G is a substrate of TbPLK in *T. brucei*. (**A**). Co-immunostaining of KIN-G-3HA and TbPLK during the cell cycle in procyclic form. Open arrowheads indicate KIN-G signal at the centrin arm. TbPLK signal at different structures is indicated. BB: basal body; CA: centrin arm; FC: flagella connector; FAZt: FAZ tip. Scale bars: 5 μm. (**B**). TbPLK phosphorylates KIN-G *in vitro*. The asterisk indicates a non-specific band. (**C**). *In vitro* phosphorylated KIN-G migrates slower than non-phosphorylated KIN-G on SDS-PAGE. (**D**). Co-immunoprecipitation of native KIN-G protein by WDR2-3HA from cells treated with or without GW843682X and from cell lysate treated with Lambda protein phosphatase (λPPase). Shown is a silver-stained 12% SDS-PAGE gel. (**E**). *In vitro* TbPLK phosphosites on KIN-G protein identified by mass spectrometry. Phosphosites highlighted in red indicate the *in vitro* and *in vivo* phosphosites. MD: motor domain; CC: coiled coil; MB: microtubule-binding motif; NB: nucleotide-binding motif. (**F**). Phosphosites within the KIN-G protein sequence spanning MB1 to MB3. Sequences highlighted in blue indicate the microtubule-binding motifs (MB1, MB2 and MB3), and sequences highlighted in green indicate the nucleotide-binding motifs (NB2 and NB3). Phosphosites highlighted in red in the yellow box indicate the *in vitro* and *in vivo* phosphosites. (**G**). Effect of GW843682X treatment on the phosphorylation levels of Thr301 and Ser569. Shown is the % of reduction of phosphorylated peptides after GW843682X treatment. Error bars indicate S.D. from three independent experiments. (**H**). Percentage of phosphosites and non-phosphosites of Thr301- and Ser569-containing peptides in wild-type trypanosome cells. Error bars indicate S.D. from three independent experiments.

We asked whether TbPLK may phosphorylate KIN-G, so we performed *in vitro* kinase assay using purified recombinant TbPLK and KIN-G proteins and ATP-γ-S for thio-phosphorylation and subsequent detection of thio-phosphorylated proteins by the anti-ThioP antibody (Allen et al., 2007). We detected auto-phosphorylation of TbPLK, indicating active TbPLK, and phosphorylation of KIN-G (Figure 1B), suggesting that KIN-G is an *in vitro* substrate for TbPLK. To identify the *in vitro* TbPLK phosphosites on KIN-G, we performed *in vitro* kinase assay with purified recombinant TbPLK and KIN-G proteins and regular ATP, and then separated the proteins by SDS-PAGE, which showed that the phosphorylated KIN-G protein migrated slower than the non-phosphorylated KIN-G protein (Figure 1C). The phosphorylated KIN-G protein band was excised from the gel and analyzed by mass spectrometry to identify the *in vitro* TbPLK phosphosites on KIN-G.

Because the dephosphorylated KIN-G migrates faster on SDS-PAGE (Zhou et al., 2025), we asked whether inhibition of TbPLK activity may cause KIN-G mobility shift on SDS-PAGE gel. To avoid the potential interference of epitope-tagging with KIN-G mobility shift, we co-immunoprecipitated native, non-tagged KIN-G by its interacting protein WDR2 (Zhou et al., 2025), which was tagged with a C-terminal triple HA epitope. We treated the trypanosome cells expressing WDR2-3HA with GW843682X, a potent TbPLK inhibitor (Li et al., 2010), and performed immunoprecipitation with the anti-HA antibody to co-immunoprecipitate KIN-G and WDR2-3HA. Co-immunoprecipitation was also performed using control cells, and the cell lysate was treated with or without Lambda protein phosphatase before immunoprecipitation. The co-immunoprecipitated KIN-G band from the Lambda protein phosphatase-treated sample and the GW843682X-treated cells appeared to migrate slightly faster than that from the non-treated cells (Figure 1D). Nonetheless, the KIN-G protein bands were excised from the silver-stained gel and analyzed by mass spectrometry to identify the *in vivo* TbPLK phosphosites.

Mass spectrometric analysis of the *in vitro* phosphorylated KIN-G protein (Figure 1C) identified 29 *in vitro* TbPLK phosphosites (Figure 1E), among which 10 sites were identified as *in vivo* phosphosites in previous work (Nett et al., 2009; Zhou et al., 2025) and 10 sites were identified as *in vivo* phosphosites in the current study from the co-immunoprecipitated KIN-G (Figure 1D). Ten out of these 20 *in vivo* phosphosites are located within the motor domain of KIN-G, with Thr273 and Thr289 located in the microtubule-binding motif #2 and Thr301 located in the microtubule-binding motif #3 (Figure 1E, F). The remaining ten sites are in the C-terminal coiled-coil motifs and in the loops between coiled-coil motifs (Figure 1E). The identification of these 20 *in vitro* TbPLK phosphosites that are also phosphorylated *in vivo* in trypanosome cells suggests that these sites could be potential *in vivo* TbPLK phosphosites. However, when the phosphosites were compared between GW843682X-treated and non-treated samples, only two phosphosites, Thr301 and Ser569, were found to be reduced after GW843682X treatment, with the Thr301 phosphosite-containing peptides and the Ser569 phosphosite-containing peptides decreased by ∼27% and 100%, respectively, after GW843682X treatment (Figure 1G). The less reduction in Thr301 phosphorylation than Ser569 phosphorylation after GW843682X treatment suggests slower dephosphorylation of Thr301 in cells after inhibition of TbPLK activity. Alternatively, another protein kinase may also phosphorylate Thr301 in the absence of TbPLK activity, causing the observed less reduction in Thr301 phosphorylation. Nonetheless, this result suggests that Thr301 and Ser569 are *in vivo* phopshosites of TbPLK.

We next analyzed the ratio of the phosphorylated/non-phosphorylated Ser301- and Ser569-containing peptides in trypanosome cells. The results showed that ∼14% of the Ser301-containing peptides and ∼28% of the Ser569-containing peptides were phospho-peptides (Figure 1H), indicating that a small proportion of the native KIN-G protein is phosphorylated at these two sites by TbPLK in trypanosome cells. Since KIN-G and TbPLK co-localize during early S-phase (Figure 1A), it is likely that KIN-G is phosphorylated during early S-phase and dephosphorylated during other cell cycle stages.

### TbPLK phosphorylation of KIN-G disrupts KIN-G binding to microtubules

Because TbPLK co-localizes with KIN-G during early cell cycle stages at the centrin arm and phosphorylates KIN-G on Thr301 in the microtubule-binding motif #3 (Figure 1), we tested the effect of TbPLK phosphorylation on the microtubule-binding activity and/or microtubule-gliding activity of KIN-G by *in vitro* microtubule-binding and -gliding assay (Zhou et al., 2024). Purified recombinant KIN-G protein was first settled onto glass coverslips in a flow chamber, and then rhodamine-labeled microtubules were added into the flow chamber, where microtubules first bound to KIN-G and then moved toward their minus-ends (opposite to the plus end-directed moving direction of KIN-G) (Zhou et al., 2024). We detected strong binding of microtubules when only KIN-G was settled on the coverslips, but we observed a significant reduction in the number of the attached microtubules when TbPLK was co-settled with KIN-G (Figure 2A, B, Figure 2-video 1, and Figure 2-video 2). Neither the microtubule-binding activity nor the microtubule-gliding activity of KIN-G was affected when it was co-settled with the kinase-dead TbPLK mutant TbPLK^K70R^ (Figure 2A-C and Figure 2-video 3). These results suggest that TbPLK activity inhibits the microtubule-binding activity of KIN-G.

**Figure 2.**
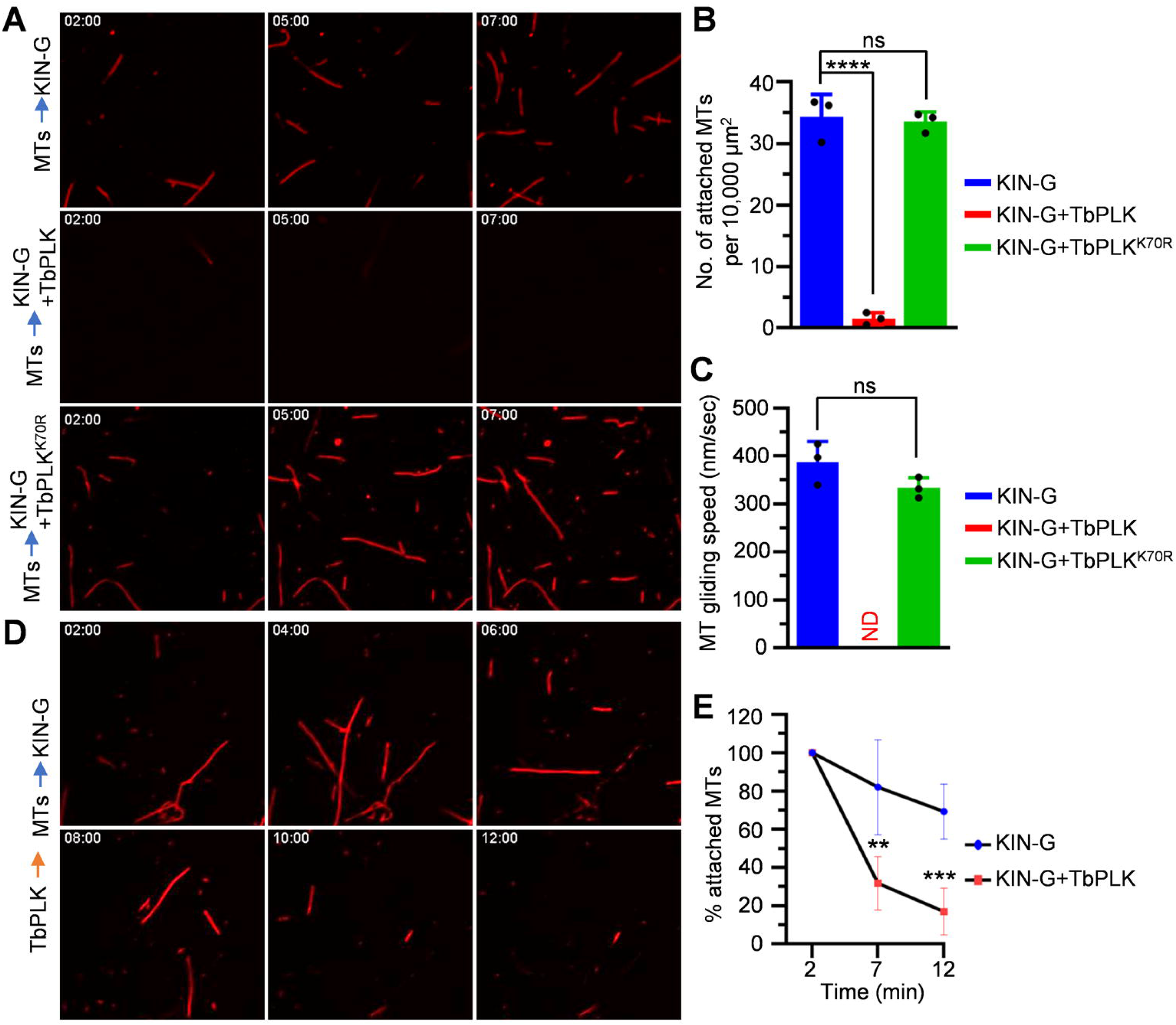
Phosphorylation of KIN-G by TbPLK disrupts the microtubule-binding activity of KIN-G. (**A**). Pre-incubation of KIN-G with TbPLK, but not TbPLK^K70R^, disrupted KIN-G microtubule-binding activity. KIN-G, KIN-G and TbPLK mixture, and KIN-G and TbPLK^K70R^ mixture were first attached to coverslips and then microtubules (MTs) were added into the chamber. (**B**). Quantitation of KIN-G-bound microtubules in the presence or absence of TbPLK or in the presence of TbPLK^K70R^. Error bars indicate S.D. from three independent experiments. ****: *p*<0.0001; ns: no significance (one-way ANOVA). (**C**). Measurement of the microtubule-gliding speed of KIN-G in the presence or absence of TbPLK or in the presence of TbPLK^K70R^. Error bars indicate S.D. from three independent experiments. ND: not done. ns: no significance (one-way ANOVA). (**D**). TbPLK disrupted KIN-G microtubule-binding activity. KIN-G was first attached to coverslips, MTs were added into the chamber, and finally TbPLK was added into the chamber. (**E**). Quantitation of KIN-G-bound microtubules following the incubation with or without TbPLK. Error bars indicate S.D. from three independent experiments. **: *p*<0.01; ***: *p*<0.001 (Student’s *t*-test).

We next performed *in vitro* microtubule-binding and -gliding assay by adding TbPLK after incubating microtubules with the settled KIN-G on the glass coverslip (Figure 2D and Figure 2-video 4). Without adding TbPLK, the number of bound microtubules decreased by an average of ∼30% after 12 min of incubation, whereas when TbPLK was added into the chamber, the number of bound microtubules decreased by an average of ∼83% after 12 min of incubation (Figure 2D, E), demonstrating that TbPLK-mediated phosphorylation caused the detachment of the microtubules that had already attached to KIN-G before the addition of TbPLK.

### Phosphorylation of Thr301 in KIN-G by TbPLK impairs the binding of KIN-G to microtubules

Because TbPLK phosphorylates the Thr301 residue within the microtubule-binding motif #3 of KIN-G (Figure 1) and TbPLK inhibits the microtubule-binding activity of KIN-G *in vitro* (Figure 2), we asked whether phosphorylation of Thr301 may disrupt the microtubule-binding activity of KIN-G *in vitro*. We mutated Thr301 to generate phospho-mimic and phospho-deficient mutants, KIN-G^T301D^ and KIN-G^T301A^, respectively, purified recombinant proteins, and used them in the *in vitro* microtubule-binding and -gliding assays. The results showed that KIN-G^T301D^ lost microtubule-binding activity, but neither the microtubule-binding activity nor the microtubule-gliding activity of KIN-G^T301A^ was different from wild-type KIN-G (Figure 3A-C, Figure 3-video 1, Figure 3-video 2, and Figure 3-video 3). These results suggest that phosphorylation of Thr301 disrupts the microtubule-binding activity of KIN-G. We also mutated the Thr284 residue, an *in vitro* TbPLK phosphosite located in the microtubule-binding motif #2 (Figure 1E), to generate phospho-mimic and phospho-deficient mutants, KIN-G^T284D^ and KIN-G^T284A^, respectively, and tested their microtubule-binding and -gliding activities. Neither KIN-G^T284D^ nor KIN-G^T284A^ showed any defects in microtubule-binding and -gliding activities (Figure 3A-C, Figure 3-video 4, and Figure 3-video 5).

**Figure 3.**
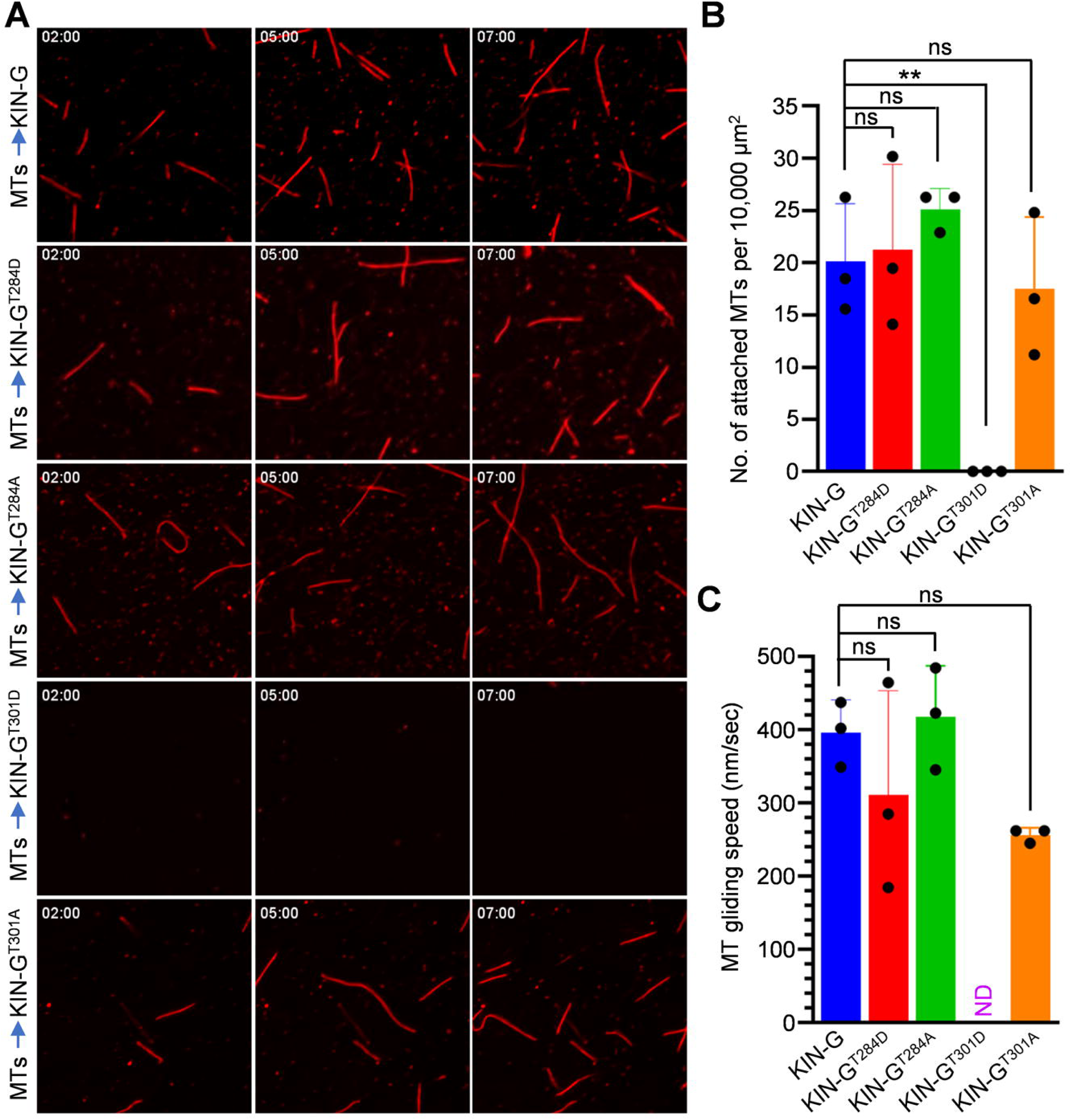
Phosphorylation of Thr301 on KIN-G by TbPLK disrupts the microtubule-binding activity of KIN-G. (**A**). Microtubule-binding activity of KIN-G and its Thr284 and Thr301 mutants. (**B**). Quantitation of the bound microtubules of KIN-G and its mutants. Error bars indicate S.D. from three independent experiments. **: *p*<0.01; ns: no significance (one-way ANOVA). (**C**). Measurement of the microtubule-gliding speed of KIN-G and its mutants. Error bars indicate S.D. from three independent experiments. ND: not done. ns: no significance (one-way ANOVA).

### Expression of Thr301 phospho-mimic mutant causes defective cell proliferation

To investigate the potential effects of T301 phosphorylation on the cellular function of KIN-G in *T. brucei*, we performed RNAi complementation experiments by ectopically expressing KIN-G^T301D^, KIN-G^T301A^, and KIN-G, which were recoded to resist RNAi and were tagged with a C-terminal triple HA epitope, in the KIN-G RNAi cell line. Immunofluorescence microscopy showed that like wild-type KIN-G, both the T301A and T301D mutants were localized to the centrin arm (Figure 4A), suggesting that phosphorylation of KIN-G does not affect its subcellular localization. Western blotting confirmed the knockdown of the endogenously PTP-tagged KIN-G and the ectopic expression of triple HA-tagged wild-type and mutant KIN-G proteins (Figure 4B). Expression of wild-type KIN-G and KIN-G^T301A^, but not KIN-G^T301D^, rescued the growth defects of KIN-G RNAi cells (Figure 4C). We quantitated the cell types of different nucleus and kinetoplast configurations in KIN-G RNAi cells and the three RNAi complementation cells. The KIN-G^T301D^ complementation cells, but not the KIN-G complementation cells and the KIN-G^T301A^ complementation cells, accumulated 2N1K (two nuclei and one kinetoplast) cells and xNyK (x>2; y≥1) cells, similar to the phenotype caused by KIN-G RNAi (Figure 4D). These results suggest that hyperphosphorylation of Thr301 disrupts the cellular function of KIN-G and that dephosphorylation of T301 is required for KIN-G to execute its function.

**Figure 4.**
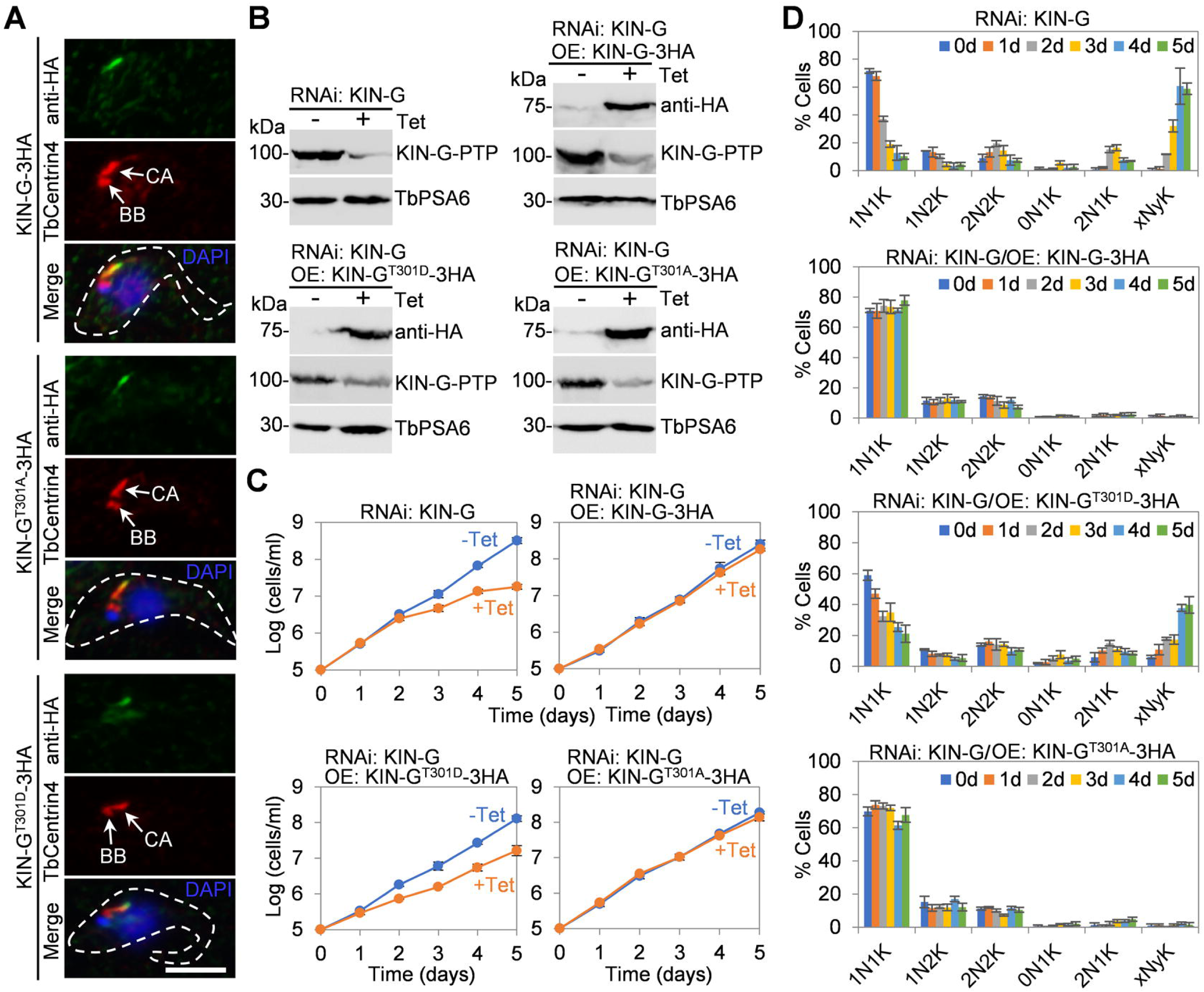
Expression of Thr301 phospho-mimic mutant of KIN-G disrupts cell proliferation. (A). Subcellular localization of ectopically expressed KIN-G, KIN-G^T301A^, KIN-G^T301D^, co-stained with TbCentrin4. BB: basal body; CA: centrin arm. Scale bar: 5 μm. (**B**). Western blotting to detect the levels of ectopically 3HA-tagged KIN-G and its mutants and the endogenously PTP-tagged KIN-G before and after tetracycline induction for 48 hours. TbPSA6 serves as loading control. (**C**). Growth curves of KIN-G RNAi cell line and its complementation cell lines expressing KIN-G, KIN-G^T301A^, or KIN-G^T301D^. OE: overexpression. Error bars indicate S.D. from three independent experiments. (**D**). Quantitation of the numbers of nuclei (N) and kinetoplasts (K) of KIN-G RNAi cell line and its complementation cell lines. 100 cells were counted for each time point, and error bars indicated S.D. from three independent experiments.

### Expression of Thr301 phospho-mimic mutant disrupts centrin arm biogenesis and Golgi duplication

We examined the centrin arm in KIN-G^T301D^ complementation cells by immunofluorescence microscopy using anti-TbCentrin4 and the pan-centrin antibody 20H5. Measurement of the length of the new and old centrin arm structures showed that many KIN-G^T301D^ complementation cells contained a shorter new centrin arm, with the average length of the new centrin arm significantly reduced after induction of KIN-G^T301D^ expression for 48 h (Figure 5A, B). The results mimicked the deficiency in the biogenesis of the new centrin arm in KIN-G RNAi cells (Zhou et al., 2024), suggesting that hyperphosphorylation of Thr301 disrupts the function of KIN-G in promoting centrin arm biogenesis.

**Figure 5.**
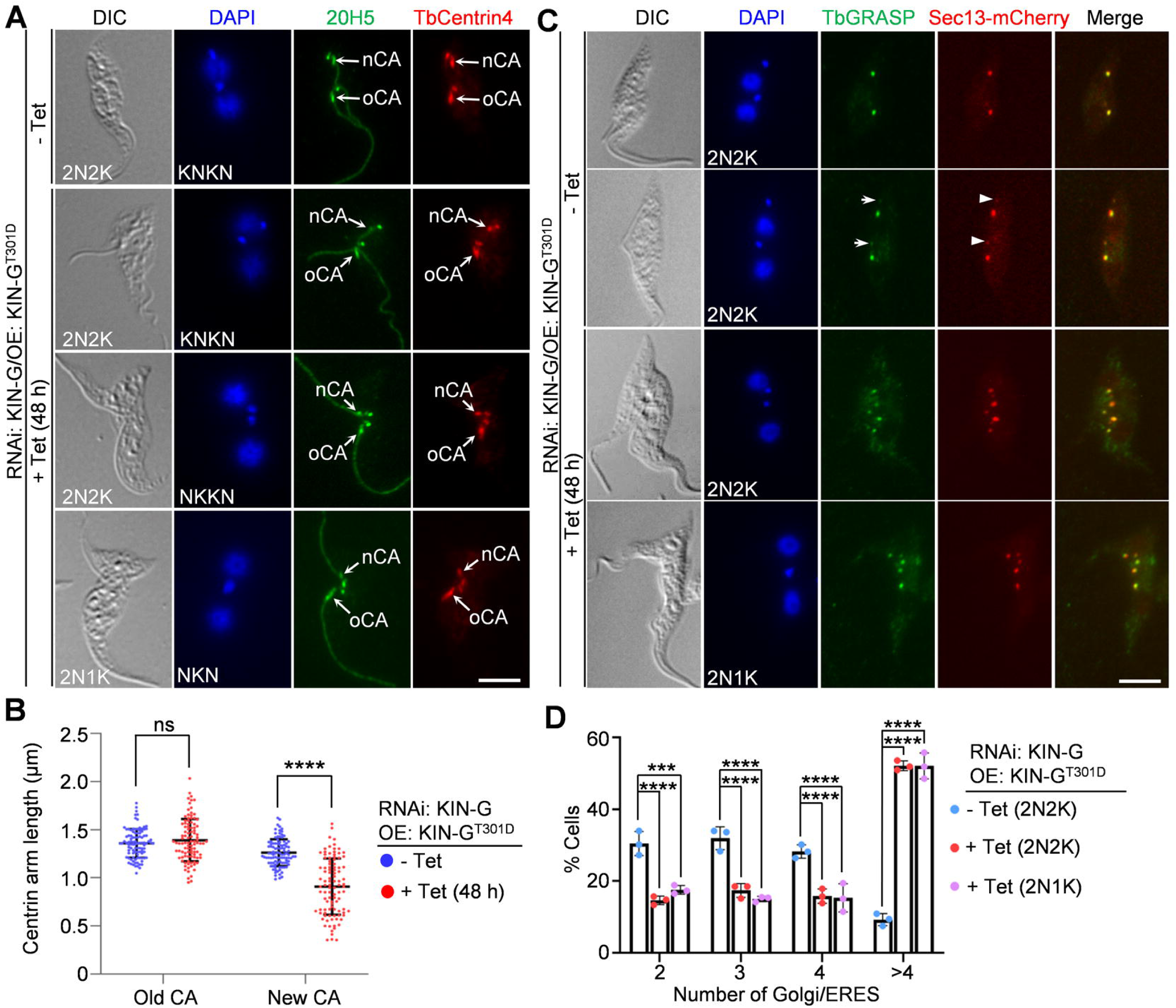
TbPLK phosphorylation of Thr301 on KIN-G disrupts centrin arm and Golgi biogenesis. (**A**). Effect of the expression of KIN-G^T301D^ on centrin arm formation. Cells were co-immunostained with the pan-centrin antibody 20H5 and the anti-TbCentrin4 antibody. Scale bar: 5 μm. (**B**). Measurement of centrin arm length in KIN-G RNAi cells expressing Thr301 phospho-mimic of KIN-G before and after tetracycline induction. 100 cells were used for measurement for each time point, and error bars indicated S.D. ns: no significance; ****: *p*<0.0001 (Student’s *t*-test). (**C**). Effect of the expression of KIN-G^T301D^ on the biogenesis of Golgi and ER exit site (ERES). Cells were co-immunostained with the anti-TbGRASP antibody to detect TbGRASP. ERES was labeled by mCherry-tagged Sec13. Scale bar: 5 μm. (**D**). Quantitation of cells with different numbers of Golgi/ERES in control and KIN-G RNAi cells. 100 cells were counted for each time point, and error bars indicated S.D. from three independent experiments. ***: *p*<0.001; ****: *p*<0.0001 (one-way ANOVA).

We next examined Golgi duplication in KIN-G^T301D^ complementation cells by immunofluorescence microscopy using the anti-TbGRASP antibody. We also endogenously epitope-tagged the COPII coat protein subunit Sec13 with mCherry to serve as a marker for the ER exit site (ERES). In non-induced control cells, two major Golgi/ERES (strong TbGRASP/Sec13 fluorescence signal) were detected in bi-nucleated cells (Figure 5C). Additionally, one or two smaller Golgi (weaker TbGRASP signal), each of which associated with a smaller ERES (weaker Sec13 signal), were also detected near the major Golgi/ERES in some bi-nucleated cells (Figure 5C, arrows and arrowheads, respectively). In the KIN-G^T301D^ complementation cells, the number of bi-nucleated cells containing two, three, or four Golgi/ERES was significantly reduced, whereas the number of bi-nucleated cells containing more than four Golgi/ERES was significantly increased (Figure 5C, D). While these results suggest that Golgi duplication was impaired in KIN-G^T301D^-expressing cells, it remains unclear whether there was any Golgi biogenesis/assembly defect, which would require ultra-structural microscopic analysis using electron microscopy or Cryo-electron tomography and is beyond the scope of this study. Although the use of conventional immunofluorescence microscopy with TbGRASP as a Golgi marker is unable to detect any structural defects of Golgi, these results suggest that hyperphosphorylation of Thr301 causes altered Golgi organization or duplication.

### Expression of Thr301 phospho-mimic mutant impairs FAZ elongation and flagellum positioning

We investigated the potential effect of expression of KIN-G^T301D^ on the elongation of the new FAZ by immunofluorescence microscopy using the anti-CC2D antibody (Zhou et al., 2011), which showed that the length of the new FAZ in the bi-nucleated KIN-G^T301D^ complementation cells was significantly shorter than that of the non-induced bi-nucleated cells (Figure 6A, B). These results suggest that Thr301 hyperphosphorylation impairs FAZ elongation. Notably, in these bi-nucleated KIN-G^T301D^ complementation cells, the size of the NFD cell was also smaller and is positively correlated to the length of the new FAZ (Figure 6C). This result confirms that the FAZ length determines the cell size in trypanosomes (Zhou et al., 2011).

**Figure 6.**
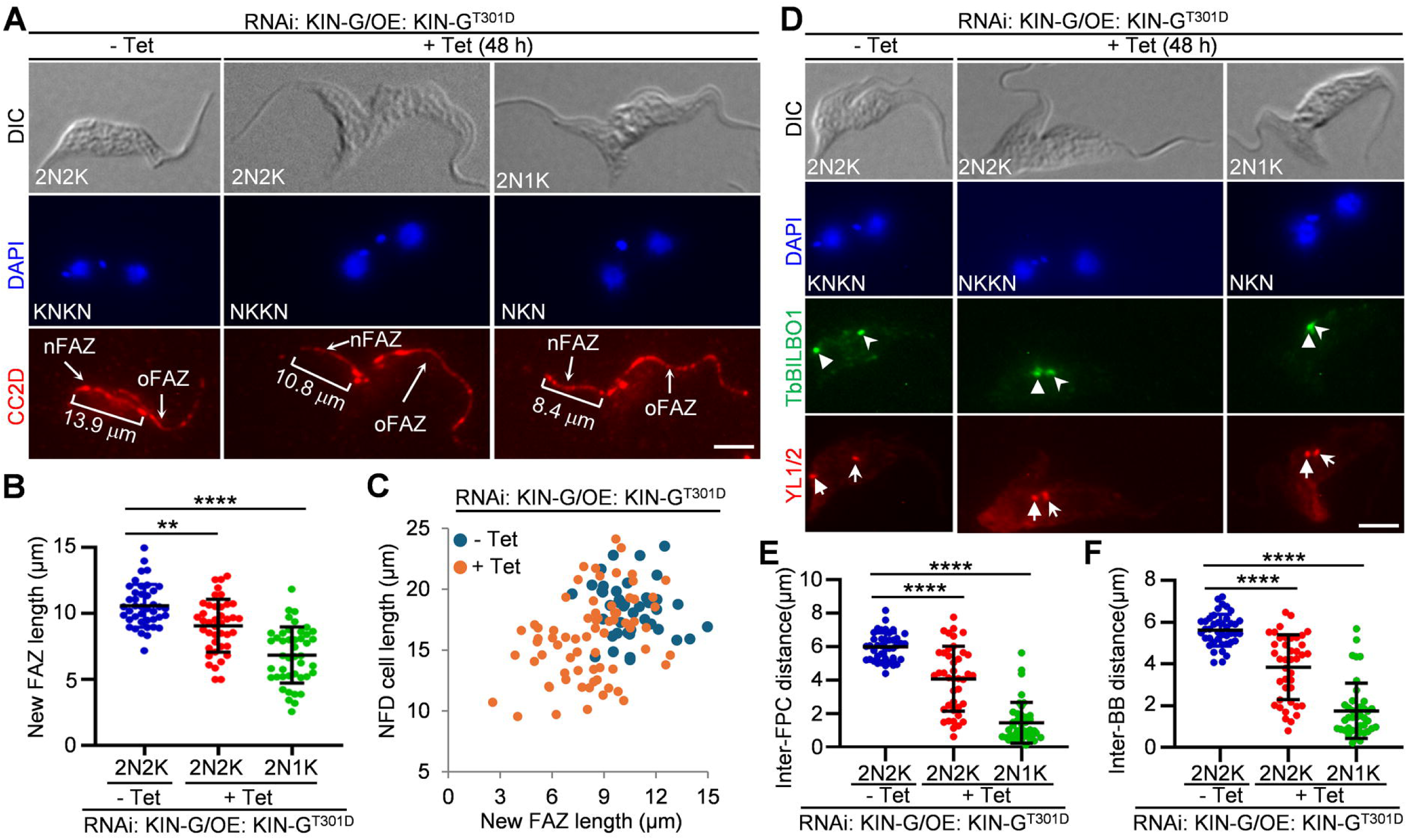
TbPLK phosphorylation of Thr301 on KIN-G impairs flagellar inheritance. (**A**). Effect of the expression of KIN-G^T301D^ on the elongation of the flagellum attachment zone (FAZ) filament. Cells were immunostained with the anti-CC2D antibody. Scale bar: 5 μm. (**B**). Measurement of the length of the new and the old FAZ filaments in control and KIN-G RNAi cells expressing KIN-G^T301D^. 100 cells were used for measurement for each time point, and error bars indicated S.D. **: *p*<0.01; ****: *p*<0.0001 (one-way ANAVA). (**C**). Measurement of the cell body length of the new-flagellum daughter (NFD) cell and its correlation with the length of the new FAZ for control cells and KIN-G RNAi cells expressing KIN-G^T301D^. 100 cells were used for measurement for each time point. (**D**). Effect of the expression of KIN-G^T301D^ on the segregation of flagellar pocket collar (FPC) and basal body (BB). Cells were co-immunostained with anti-TbBILBO1 antibody and YL1/2 antibody. Scale bar: 5 μm. (**E, F**). Measurement of the inter-FPC distance (**E**) and inter-BB distance (**F**) in control cells and KIN-G RNAi cells expressing Thr301 phospho-mimic mutant of KIN-G. 100 cells were used for measurement for each time point, and error bars indicated S.D. ****: *p*<0.0001 (one-way ANOVA).

We next examined the segregation of the flagellum-associated cytoskeletal structures in KIN-G^T301D^ complementation cells by co-immunofluorescence microscopy with YL1/2, which labels the mature basal body (Woods et al., 1989), and the anti-TbBILBO1 antibody, which labels the flagellar pocket collar (Bonhivers et al., 2008). In the bi-nucleated KIN-G^T301D^ complementation cells, the duplicated basal bodies/flagellar pocket collars were not far segregated (Figure 6D), with the average distance between the duplicated structures reduced significantly after tetracycline induction for 48 h (Figure 6E, F). These results suggest that hyperphosphorylation of Thr301 inhibits the segregation of the flagellum-associated cytoskeletal structures. Because the flagellum is nucleated from the mature basal body, these results suggest that hyperphosphorylation of Thr301 disrupts flagellum positioning.

### Expression of Thr301 phospho-mimic mutant impairs cell division plane placement

Previous work showed that knockdown of KIN-G by RNAi caused cytokinesis defects (Zhou et al., 2024), likely due to the mis-placement of the cell division plane, but this phenotype was not explored in that study. In procyclic trypanosomes, the cell division plane is placed along the longitudinal cell axis from the anterior tip of the NFD cell to the nascent posterior of the OFD cell (Wheeler et al., 2013). To investigate the potential defect in cell division plane placement in KIN-G RNAi cells, we used the orphan kinesin protein KLIF as a marker for the cell division plane (Zhou et al., 2018a). In the non-induced control cells, the leading edge of the KLIF-labeled cell division plane was always placed at the mid-portion of the NFD cell (Figure 7A, yellow arrowhead). In contrast, in the dividing KIN-G RNAi cells, the leading edge of the KLIF-labeled cell division plane was placed near the posterior cell tip (Figure 7A, yellow arrowheads). To further confirm this defect, we investigated the positioning of the nascent posterior of the OFD cell in KIN-G RNAi cells. The nascent posterior is formed near the mid-portion of the NFD cell through microtubule bundling and cytoskeleton remodeling during late stages of the cell cycle (Wheeler et al., 2013). Thus, the NFD cell inherits the old, existing cell posterior, whereas the OFD cell inherits the newly formed or nascent cell posterior. We used the epitope-tagged GB4 protein, a microtubule plus end-localized protein (Rindisbacher et al., 1993), as a marker to label the cell posteriors (Figure 7B). In control cells, the GB4-labeled nascent posterior of the OFD cell was located at the mid-portion of the NFD cell, whereas in KIN-G RNAi cells, the GB4-labeled nascent posterior of the OFD cell was placed next to the GB4-labeled existing posterior of the NFD cell (Figure 7B). This defect in nascent posterior positioning was confirmed by measurement of the inter-posterior distance, which showed a significant reduction in KIN-G RNAi cells (Figure 7C).

**Figure 7.**
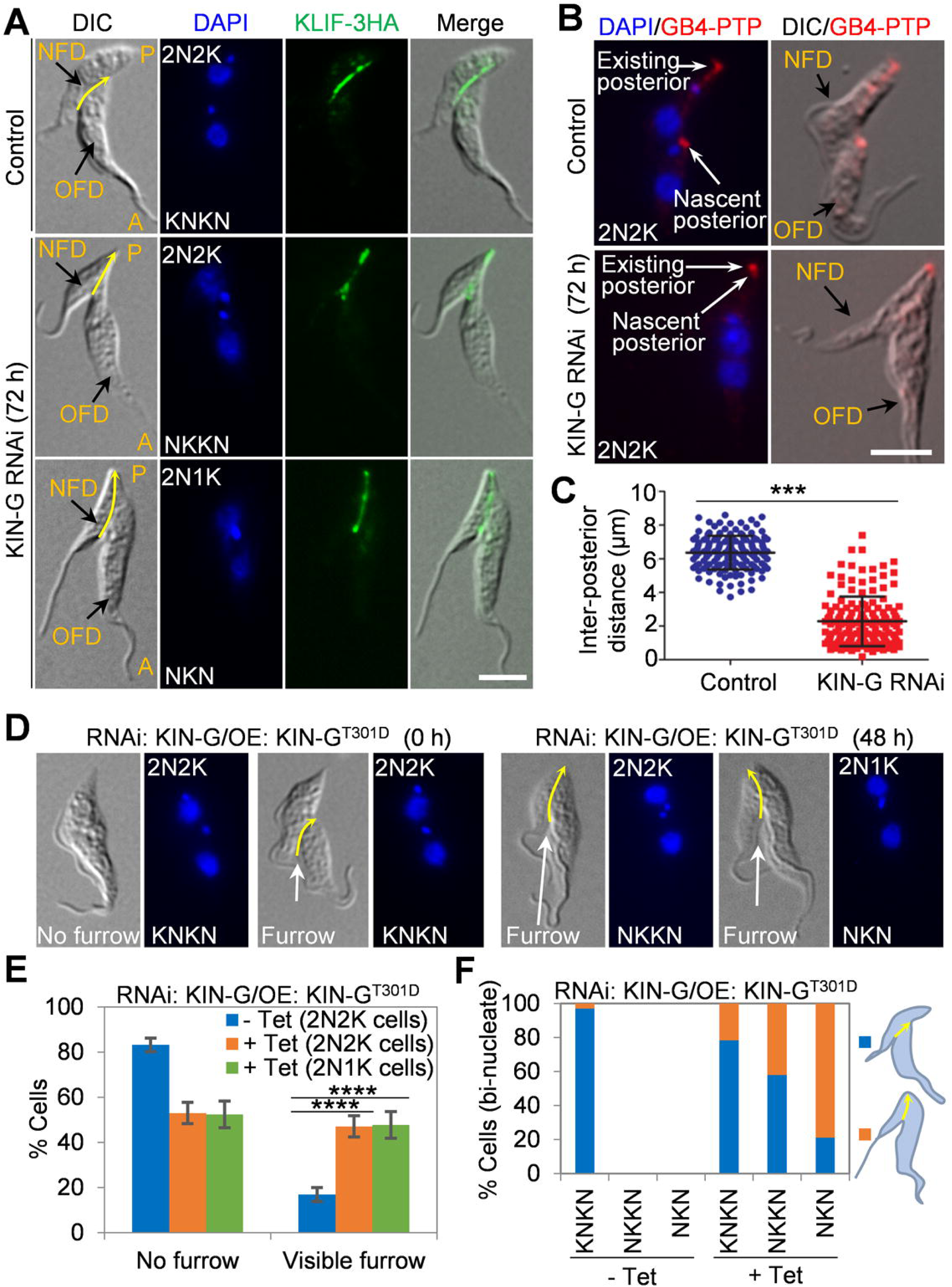
Phosphorylation of KIN-G at Thr301 by TbPLK disrupts cell division plane placement. (**A**). Immunofluorescence microscopy to detect the cell division plane with endogenous triple HA-tagged KLIF in dividing cells from non-induced control and KIN-G RNAi cells. Yellow arrows indicate the KLIF-marked cell division plane. NFD: new-flagellum daughter; OFD: old-flagellum daughter; P: posterior; A: anterior. Scale bar: 5 μm. (**B**). Immunofluorescence microscopy to detect the NFD posterior and the OFD nascent posterior with PTP-tagged GB4 protein. Scale bar: 5 μm. (**C**). Measurement of the inter-posterior distance of bi-nucleated cells from non-induced control and KIN-G RNAi cells. 100 cells were used for measurement for each time point, and error bars indicated S.D. ***, *p*<0.001 (Student’s *t*-test). (**D**). Non-induced and tetracycline-induced KIN-G RNAi cells expressing KIN-G^T301D^. Shown are a non-dividing cell without a visible cleavage furrow and three dividing cells with a visible cleavage furrow. Yellow arrows indicate the cell division plane. Scale bar: 5 μm. (**E**). Quantitation of bi-nucleated cells with or without a visible cleavage furrow from non-induced and tetracycline-induced KIN-G RNAi cells expressing KIN-G^T301D^. 100 cells were counted for each time point, and error bars indicated S.D. from three independent experiments (n=3). ****, *p*<0.0001 (one-way ANOVA). (**F**). Percentage of dividing bi-nucleated cells with a normally placed cell division plane or an abnormally placed cell division plane from non-induced and tetracycline-induced KIN-G RNAi cells expressing KIN-G^T301D^. 100 cells were used for measurement for each time point.

The KIN-G^T301D^ complementation cells appeared to have similar cytokinesis defects (Figure 4C, D). The number of bi-nucleated cells undergoing cytokinesis or with a visible cleavage furrow was significantly increased upon tetracycline induction (Figure 7D, E), suggesting that the KIN-G^T301D^ complementation cells were able to initiate cytokinesis, but failed to complete cytokinesis. Notably, those KIN-G^T301D^ complementation cells with a visible cleavage furrow contained a mis-placed cell division plane (Figure 7D, yellow arrows). Notably, mis-placement of the cell division plane was only observed in the bi-nucleated cells with the NKKN configuration and the NKN configuration, i.e., cells with defective kinetoplast segregation (Figure 7F). Together, these results suggest that hyperphosphorylation of Thr301 disrupts cell division plane placement, mimicking the defect caused by KIN-G RNAi.

## Discussion

In this report, we identified the orphan kinesin KIN-G as a substrate of TbPLK, one of the two Polo-like kinase homologs in *T. brucei* (Kurasawa et al., 2020), and discovered a negative role of TbPLK phosphorylation in regulating the biochemical activity and the cellular function of KIN-G in the procyclic form of *T. brucei*. One of the intriguing questions is how TbPLK-phosphorylation of KIN-G negatively regulates the function of KIN-G. KIN-G is a microtubule plus end-directed motor protein, and its motor activity is required for cellular function (Zhou et al., 2024), suggesting that KIN-G needs to travel along microtubules to execute its function. The best candidate microtubules that KIN-G may associate with are the specialized set of four microtubules (MtQ), which have their plus ends directed toward the distal tip of the cell in *T. brucei* (Gull, 1999; Robinson et al., 1995). Presumably, KIN-G may be first recruited to the centrin arm through binding to WDR2 (Zhou et al., 2025) and then it associates with and travels along the MtQ to transport cargos in the centrin arm region to facilitate centrin arm biogenesis and Golgi duplication (Fig. 8). In this regard, the T301-phosphorylated KIN-G is unable to bind to MtQ due to the loss of microtubule-binding activity (Figures 2 and 3) and, hence, is unable to transport cargos to their destination. The lack of rescue of KIN-G RNAi by the expression of the phospho-mimic mutant KIN-G^T301D^ and the complete rescue of KIN-G RNAi by the expression of the phospho-deficient mutant KIN-G^T301A^ (Figure 4) support this notion. Additionally, these findings also suggest that dephosphorylation of Thr301, but not phosphorylation of Thr301, is required for the cellular function of KIN-G. This negative regulation of KIN-G function by TbPLK phosphorylation resembles the regulation of TbCentrin2 by TbPLK phosphorylation (de Graffenried et al., 2013), because dephosphorylation of TbCentrin2 was also found to be required for TbCentrin2 function. To dephosphorylate KIN-G and TbCentrin2, the best protein phosphatase candidate is the centrin arm-localized KPP1 (Zhou et al., 2018b), which also attenuates TbPLK kinase activity by dephosphorylating TbPLK at Thr125 (An et al., 2021). However, whether KPP1 is responsible for the dephosphorylation of KIN-G and TbCentrin2 to antagonize TbPLK-mediated phosphorylation of these two proteins remains to be explored.

**Figure 8.**
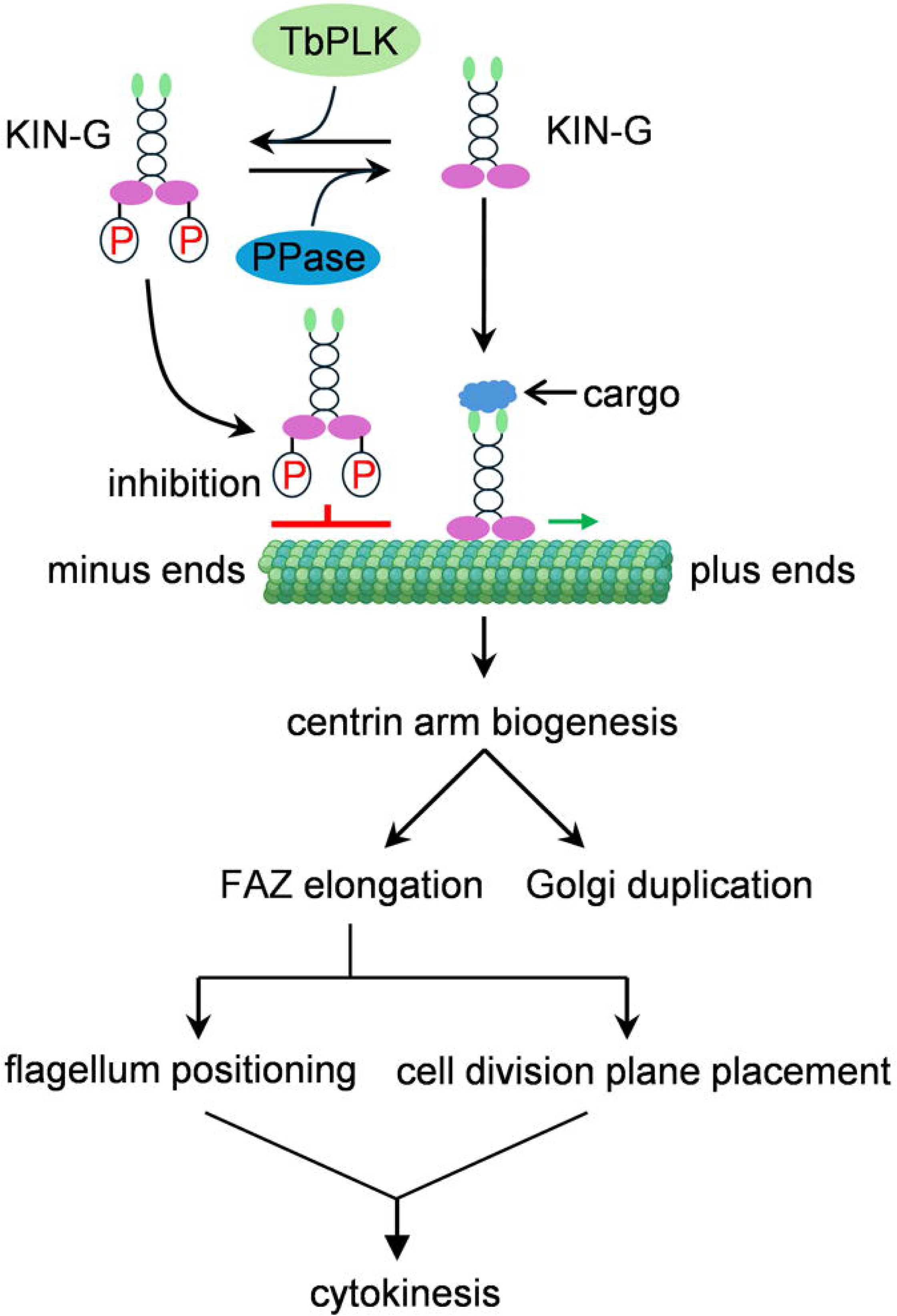
Model of KIN-G’s biochemical and cellular function and its regulation through phosphorylation by TbPLK and dephosphorylation by an unknown protein phosphatase. KIN-G is depicted as a dimer and transports cargos along the MtQ near the centrin arm region to regulate centrin arm biogenesis, which impacts Golgi duplication and FAZ elongation, the latter of which promotes flagellum positioning and cell division plane placement, thereby facilitating cytokinesis. The green arrow indicates the direction of movement of KIN-G. Phosphorylation of KIN-G by TbPLK inhibits its microtubule-binding activity, thereby disrupting KIN-G function.

Expression of the Thr301 phospho-mimic mutant in procyclic trypanosomes disrupted centrin arm formation, Golgi duplication, FAZ elongation, flagellum positioning, cell division plane placement, and cytokinesis (Figures 4-7), mimicking the defects caused by KIN-G RNAi (Zhou et al., 2024). There is an intriguing question of why KIN-G appears to regulate so many cellular processes, while it only localizes to the centrin arm throughout the cell cycle (Zhou et al., 2024). Previous work has demonstrated that the defect in centrin arm integrity impairs FAZ elongation, flagellum positioning, cell division plane placement, and Golgi duplication (de Graffenried et al., 2013; de Graffenried et al., 2008; He et al., 2005; Zhou et al., 2010; Zhou et al., 2025; Zhou et al., 2024). In this regard, all the defects, except for defective centrin arm biogenesis, observed in KIN-G^T301D^ complementation cells and KIN-G RNAi cells are secondary effects attributed to the defective centrin arm biogenesis. Therefore, the primary function of KIN-G is to regulate centrin arm biogenesis. Because phosphorylation of Thr301 by TbPLK disrupts the microtubule-binding activity of KIN-G and, hence, its plus end-directed motility, it suggests that regulation of centrin arm biogenesis by KIN-G requires its motor activity, in agreement with the results obtained with the expression of the motility-dead mutant KIN-G^G266A^, which caused similar defects (Zhou et al., 2024).

How does the defect in centrin arm integrity impair Golgi duplication, FAZ elongation, flagellum positioning, and cell division plane placement? The Golgi apparatus in trypanosomes is located in close proximity to the centrin arm (He et al., 2005), whereas the proximal end of the FAZ is embedded within the hook complex between the centrin arm and the shank part of the hook complex (Morriswood et al., 2013). Golgi duplication is coupled to centrin arm assembly (He et al., 2005), and assembly of the FAZ occurs from its proximal end (Sunter et al., 2015; Zhou et al., 2015). Therefore, the primary role of the centrin arm likely is to act as a hub to coordinate FAZ assembly (Zhou et al., 2010) and determine the Golgi assembly site (He et al., 2005) (Figure 8). The defect in FAZ elongation, observed in KIN-G RNAi cells (Zhou et al., 2024) and KIN-G^T301D^ complementation cells (Figure 7), likely contributes to flagellum mis-positioning and cell division plane placement (Figure 8), because previous work has demonstrated that flagellum positioning relies on the faithful duplication/segregation of its associated cytoskeletal structures, including the FAZ (Ikeda and de Graffenried, 2012; Zhou et al., 2011) and that the length of the new FAZ determines the cell division plane (Zhou et al., 2011). Consequently, mis-positioning of the newly synthesized flagellum and mis-placement of the cell division plane, observed in KIN-G RNAi cells and KIN-G^T301D^ complementation cells, collectively contribute to defective cytokinesis (Figure 8).

The complete rescue of KIN-G RNAi phenotype by expression of the phospho-deficient mutant KIN-G^T301A^ (Figure 4) raised an interesting question about the role of TbPLK-mediated phosphorylation of KIN-G on this residue in trypanosome cells. Since KIN-G co-localizes with TbPLK from G1 phase to early S-phase, it is possible that KIN-G is phosphorylated during these stages and then dephosphorylated from late S-phase when TbPLK leaves the centrin arm (Figure 1A). Indeed, Thr301-phosphorylated KIN-G accounts for ∼14% of the total KIN-G population in asynchronous trypanosome cells (Figure 1H), suggesting that phosphorylation and dephosphorylation of KIN-G are controlled events during the cell cycle. However, the lack of phenotype of the KIN-G^T301A^ complementation cells and the strong phenotype of the KIN-G^T301D^ complementation cells (Figures 4-7) suggest that phosphorylation of Thr301 may act as a negative regulatory switch, but not constitutively required under normal growth conditions. Phosphorylation of Thr301 may be transient, spatially restricted, or condition-specific, so expression of the phospho-deficient T301A mutant does not disrupt the basal function. This phosphorylation may become essential under other conditions, such as stress conditions. Thus, expression of the KIN-G^T301D^ mutant, however, generates a constitutive negative regulation, and the strong phenotype caused by its expression (Figures 4-7) suggests that it is the Thr301 hyperphosphorylation that caused the defects, and that dephosphorylation of Thr301 is necessary for KIN-G function. Given the essential involvement of centrin arm biogenesis in various cellular processes, FAZ elongation, Golgi duplication, flagellum positioning, and cell division plane placement (Figures 5-7), Thr301 phosphorylation may serve as a checkpoint mechanism to prevent premature or erroneous centrin arm biogenesis, in the event when cells detect any centrin arm biogenesis errors or the unavailability of certain key centrin arm component(s). In this scenario, TbPLK phosphorylates Thr301 to temporally inactivate KIN-G, allowing cells the time to repair the error or supply the component(s). This regulation scheme is conceptually analogous to the spindle assembly checkpoint mechanism, which monitors spindle microtubule-kinetochore attachment errors to prevent premature, erroneous chromosome segregation. Nonetheless, Thr301 phosphorylation is likely a regulatory inactivation switch that is important for proper temporal or spatial control, but not required for basal viability under normal growth conditions.

TbPLK also phosphorylates KIN-G at Ser569, which is located in coiled-coil motif #3 (Figure 1E). Since the C-terminal tail in kinesin proteins is known to bind cargo or cargo adaptor proteins (Hirokawa et al., 1998), the phosphorylation of Ser569 may impact the binding of KIN-G to its cargo or cargo adaptor proteins. Future experiments will be directed to explore the potential biochemical and physiological roles of this phosphosite, which may uncover other roles of TbPLK-mediated phosphorylation of KIN-G.

## Materials and Methods

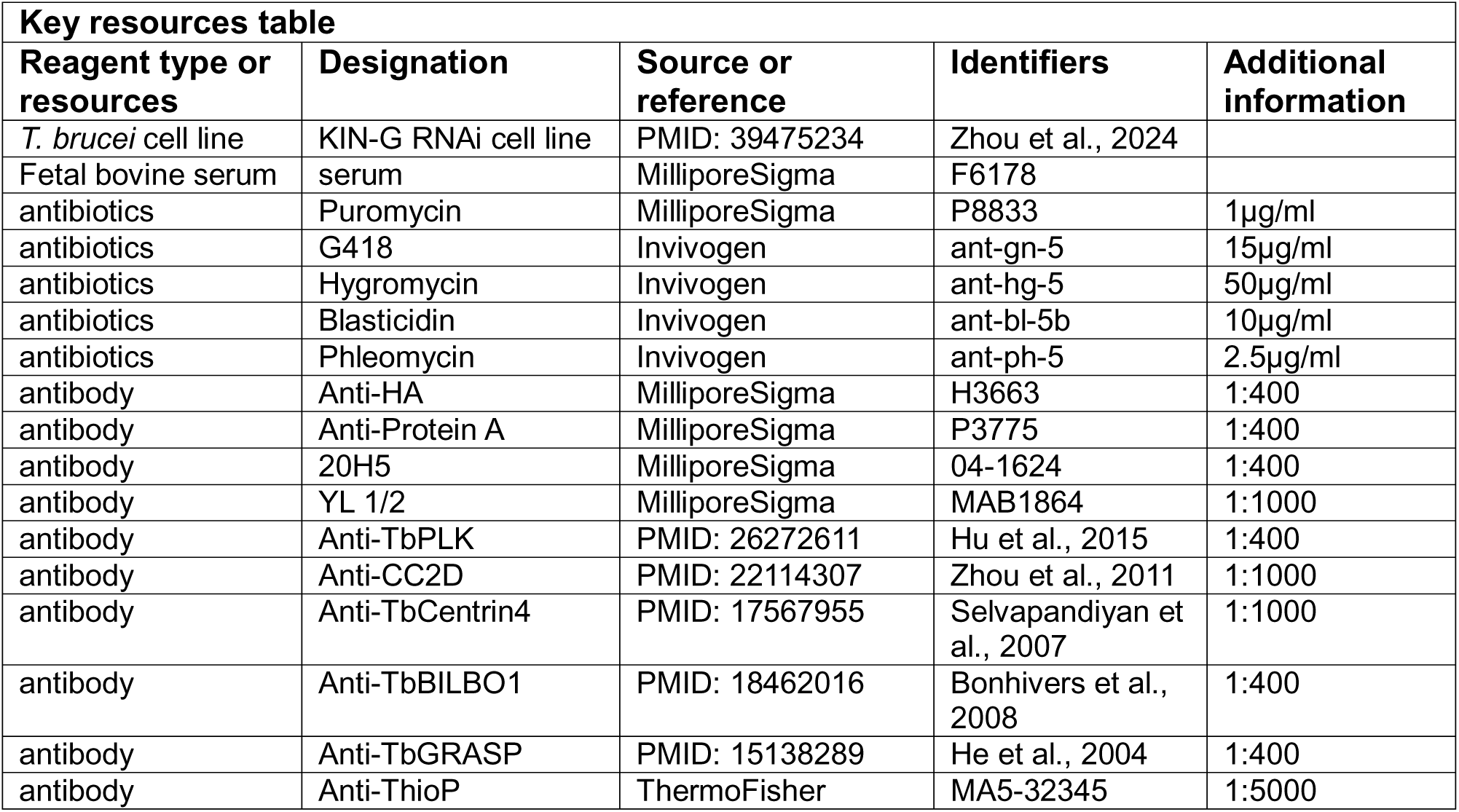
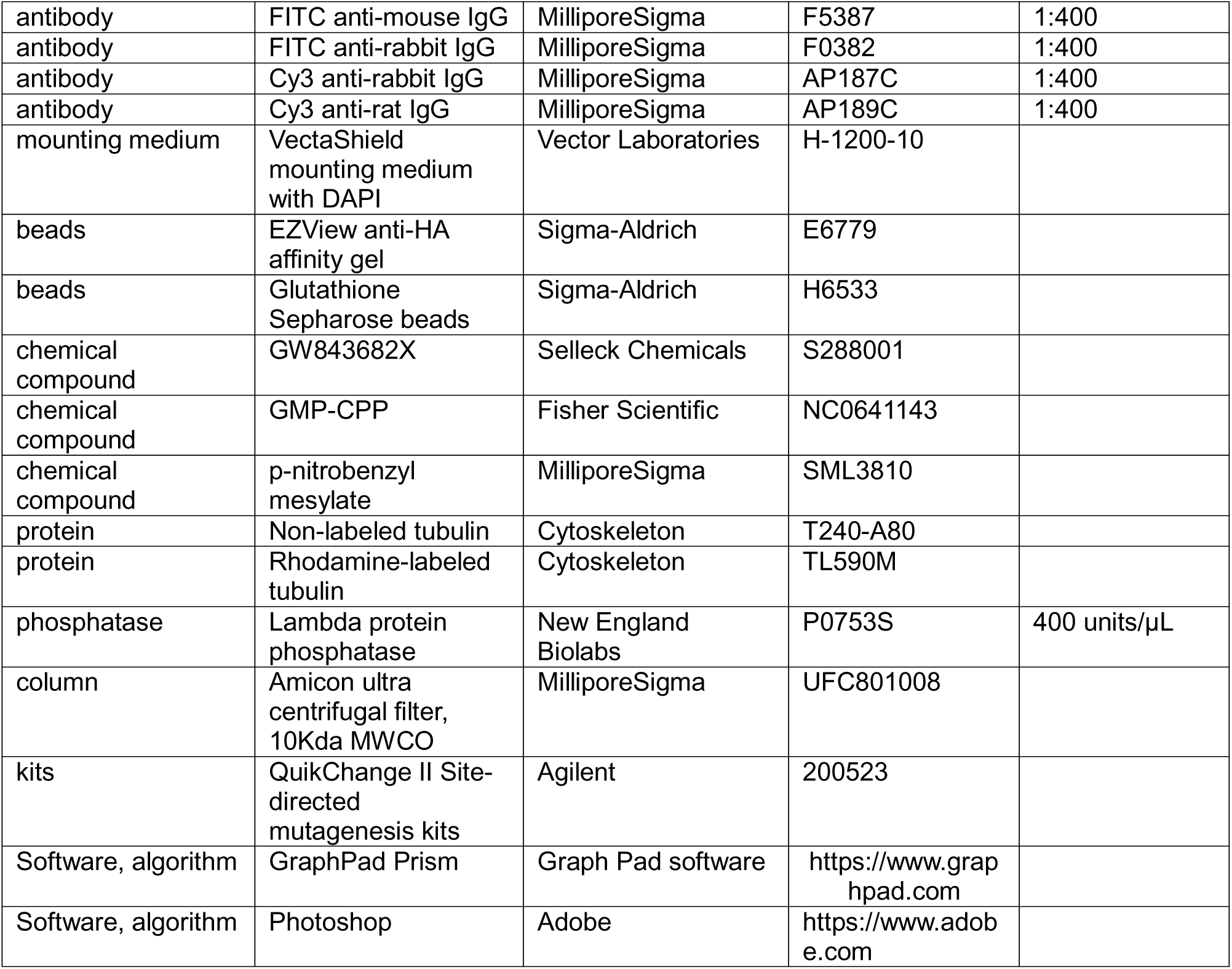

### Trypanosome cell culture, RNAi, and complementation

The procyclic form of *T. brucei* strain Lister427 was cultured in the SDM-79 medium at 27°C. The KIN-G RNAi cell line, which was generated previously (Zhou et al., 2024), was cultured in SDM-79 medium supplemented with 10% heat-inactivated fetal bovine serum (MillipreSigma, Cat# F6178), 15 µg/ml G418 (Invivogen, Cat# ant-gn-5), 50 µg/ml hygromycin (Invivogen, CAT# ant-hg-5), and 2.5 µg/ml phleomycin (Invivogen, Cat# ant-ph-5) at 27°C. Cells were diluted with fresh medium whenever the cell density reached 5×10^6^/ml. RNAi was induced with 1 µg/ml tetracycline, and cell growth was monitored daily with a hemacytometer.

The coding sequence of KIN-G was recoded to make it RNAi resistant, and the recoded KIN-G gene was cloned into pLew100-3HA-PAC vector. The pLew100-KIN-G-3HA-PAC plasmid was then used for site-directed mutagenesis to mutate the TbPLK-phosphorylated residues to aspartate or alanine, and then the resulting plasmids were used to transfect the KIN-G RNAi cell line. Transfectants were selected with 1 μg/ml puromycin (MilliporeSigma, Cat# P8833), in addition to 15 µg/ml G418, and 50 µg/ml hygromycin, and 2.5 µg/ml phleomycin, and then cloned by limiting dilution in a 96-well plate containing SDM-79 medium supplemented with appropriate antibiotics and 20% fetal bovine serum. Expression of recoded KIN-G and its phospho-mimic and phospho-deficient mutants, and knockdown of endogenous KIN-G were induced with 1 µg/ml tetracycline. Cell growth was monitored by daily counting of the cells with a hemacytometer.

### *In situ* epitope tagging of proteins

Epitope tagging of KIN-G, KLIF, GB4, and Sec13 from their respective endogenous locus was carried out using the PCR-based method (Shen et al., 2001). KIN-G was tagged with a C-terminal triple HA epitope or PTP epitope, KLIF was tagged with a C-terminal triple HA epitope, GB4 was tagged with a C-terminal PTP epitope, and Sec13 was tagged with an N-terminal mCherry tag. Transfectants were selected with appropriate antibiotics, and clonal cell lines were obtained by limiting dilution in a 96-well plate as described above.

### Immunofluorescence microscopy

To prepare intact trypanosome cells for immunofluorescence microscopy, cells were washed once with PBS, settled onto glass coverslips, fixed with methanol at −20°C for 30 min, and rehydrated with PBS at room temperature. To prepare trypanosome cytoskeletons for immunofluorescence microscopy, cells were washed once with PBS, adhered onto glass coverslips, treated with 1% Nonidet-P40 in PEME buffer (100 mM PIPES, pH6.9, 2 mM EGTA, 1 mM MgSO_4_, and 0.1 mM EDTA) for 2 seconds at room temperature, and then fixed with cold methanol at −20°C for 30 min. Intact cells or cytoskeletons on the coverslips were incubated with the blocking buffer (3% BSA in PBS) at room temperature for 1 h, and then incubated with the primary antibody at room temperature for 1 h. The following primary antibodies were used: anti-HA monoclonal antibody (MilliporeSigma, Cat# H3663, 1:400 dilution), anti-Protein A polyclonal antibody (MilliporeSigma, Cat# P3775, 1:400 dilution), anti-TbPLK antibody (1:400 dilution) (Hu et al., 2015), anti-CC2D polyclonal antibody (1:1,000 dilution) (Zhou et al., 2011), 20H5 monoclonal antibody (MilliporeSigma, Cat# 04-1624, 1:400 dilution) (He et al., 2005), YL 1/2 monoclonal antibody (MilliporeSigma, Cat# MAB1864, 1: 1000 dilution) (Woods et al., 1989), anti-TbCentrin4/LdCentrin1 polyclonal antibody (1:1000 dilution) (Selvapandiyan et al., 2007), anti-TbBILBO1 antibody (1: 400 dilution) (Bonhivers et al., 2008), and anti-TbGRASP polyclonal antibody (1:400 dilution) (He et al., 2004). Cells or cytoskeletons on the glass coverslips were washed three times (5 min each) with PBS and incubated with secondary antibodies at room temperature for 1 h. The following secondary antibodies were used: FITC-conjugated anti-mouse IgG, FITC-conjugated anti-rabbit IgG, Cy3-conjugated anti-rabbit IgG, and Cy3-conjugated anti-rat IgG (all from MilliporeSigma). Cells or cytoskeletons were washed three times (5 min each) with PBS, mounted with DAPI-containing VectaShield mounting medium (Vector Labs), and imaged with the Olympus IX71 fluorescence microscope. Images were acquired and processed using the Slidebook 5 software.

### Expression and purification of recombinant proteins

To express and purify recombinant proteins for *in vitro* microtubule-binding and gliding assays, KIN-G was cloned into the pET26 vector, and the resulting plasmid was used to generate the phospho-deficient and phospho-mimic mutants by site-directed mutagenesis. The full-length TbPLK was cloned into the pET41a vector, and the resulting plasmid was used to generate the kinase-dead mutant TbPLK^K70R^ by site-directed mutagenesis. These plasmids were each used to transform the *E. coli* BL21 strain. Expression of recombinant proteins was induced with 0.1 mM isopropyl β-d-thio-galactopyranoside at room temperature for KIN-G-6His and GST-TbPLK-6His and at 15°C overnight for GST-TbPLK^K70R^-6His. Bacterial cells expressing KIN-G-6His and its mutants, GST-TbPLK-6His, and GST-TbPLK^K70R^-6His were lysed by sonication in bacteria lysis buffer (50 mM NaH_2_PO_4_, 300 mM NaCl, 10 mM imidazole, pH 8.0), and cell lysate was cleared by centrifugation at 20,000 ×*g* for 10 min at 4°C. Cleared cell lysate was incubated with the Chelating Sepharose Fast Flow (GE Healthcare) beads charged with nickel ion at 4°C for 30 min. Beads were washed five times with 1 ml wash buffer (50 mM NaH_2_PO_4_, 1 M NaCl, 120 mM imidazole, pH 8.0) for each wash. Recombinant proteins were eluted with elution buffer (50 mM NaH_2_PO_4_, 300 mM NaCl, 250 mM imidazole, pH 8.0), and eluted proteins were concentrated and buffer-exchanged to the kinase assay buffer (10 mM HEPES, pH 7.5, 10 mM MgCl2, 50 mM NaCl, 1 mM EGTA, 1 mM DTT) with Amicon Ultra Centrifugal Filters 10K (MilliporeSigma, Cat# UFC801008).

### *In vitro* kinase assay using the thiophosphorylation method

*In vitro* kinase assay was performed by using a semisynthetic epitope to detect thiophosphorylated kinase substrates (Allen et al., 2007). Purified recombinant proteins were incubated in the kinase assay buffer (10 mM HEPES pH7.5, 10 mM MgCl_2_, 50 mM NaCl, and 0.4 mM DTT) supplemented with 1 mM ATP-γ-S at room temperature for 30 min. 50 mM p-nitrobenzyl mesylate (PNBM), which alkylates thiophosphates to form thiophosphate ester epitopes that can be recognized by the anti-thiophosphate ester antibody, was added to the kinase reaction and was incubated at room temperature for 60 min. Thiophosphorylated proteins were separated by SDS-PAGE, transferred onto a PVDF membrane, and then immunoblotted with the anti-thiophosphate ester (anti-ThioP) monoclonal antibody (ThermoFisher, Cat# MA5-32345 1:5,000 dilution,).

### Identification of *in vitro* TbPLK-phosphosites on KIN-G by mass spectrometry

Purified recombinant KIN-G-6His was mixed with purified recombinant GST-TbPLK-6His in the kinase assay buffer (see above) containing 0.2 mM ATP at room temperature for 30 min. Thiophosphorylated proteins were separated by SDS-PAGE and stained with coomassie blue dye. The gel slice containing the phosphorylated KIN-G-6His band was excised, and incubated with 160 ng trypsin at 37°C for 4 h, according to published procedures (Rosenfeld et al., 1992). Peptides were extracted with 50 ml of 50% acetonitrile and 5% formic acid, dried using SpeedVac, resuspended in 2% acetonitrile and 0.1% formic acid, and injected onto Thermo LTQ Orbitrap XL (ThermoFisher Scientific), according to published procedures (Lee and Li, 2021). Peptide samples were analyzed on an LTQ Orbitrap XL interfaced with an Eksigent nano-LC 2D plus ChipLC system (Eksigent Technologies). Samples were loaded onto a ChromXP C18-CL trap column (200 mm i.d. x 0.5 mm length) at a flow rate of 3 nL/min. Reverse-phase C18 chromatographic separation of peptides was carried on a ChromXP C18-CL column (75 mm i.d x 10 cm length) at 300 nL/min. The LTQ Orbitrap was operated in a data-dependent mode to simultaneously measure full-scan MS spectra in the Orbitrap and the five most intense ions in the LTQ by CID, respectively. In each cycle, MS1 was acquired at a target value of 1E6 with a resolution of 100,000 (m/z 400) followed by top five MS2 scan at a target value of 3E4. The mass spectrometric setting was as follows: spray voltage at 1.6 KV, and charge state screening and rejection of singly charged ion enabled. Ion selection thresholds were set at 8,000 for MS2, 35% normalized collision energy, activation Q was set at 0.25, and dynamic exclusion was employed for 30 sec. Raw data files were processed and searched against the *T. brucei* proteome database using the Mascot and Sequest HT (version 13) search engines. The search conditions were set as follows: peptide tolerance of 10 p.p.m. and MS/MS tolerance of 0.8 Da, with two missed cleavages permitted and the enzyme set as trypsin.

### Identification of *in vivo* TbPLK phosphosites on KIN-G

To identify the *in vivo* TbPLK phosphosites on KIN-G, we treated trypanosome cells (1×10^9^) expressing WDR2-3HA (for co-immunoprecipitation of native, non-tagged KIN-G) with or without 1 μM GW843682X (Selleck Chemicals, Cat# S288001) for 16 h. Cells were lysed in lysis buffer (25 mM Tris, pH 7.4, 100 mM NaCl, 1 mM DTT, 0.07% NP-40, 5% glycerol), and the lysate was cleared by centrifugation at 20,000 × *g* for 5 min at 4°C. The supernatant was incubated with 10 μl of EZview™ Red Anti-HA Affinity Gel (MilliporeSigma, Cat# E6779) for 50 min at 4°C. Beads were washed four times with lysis buffer, and then incubated with 200 units of lambda protein phosphatase (New England Biolabs, Cat# P0753S) in the phosphatase buffer (50 mM HEPES (pH 7.5), 100 mM NaCl, 2 mM DTT, 0.01% Brij 35, and 1 mM MnCl) for 20 min at 30°C. Beads were washed again and then boiled in sample buffer (50 mM Tris, pH 6.8, 1% SDS, 10% glycerol, 1% β-mercaptoethanol, 0.006% Bromophenol blue). Samples were separated by SDS-PAGE, and the gel was silver-stained as follows. The SDS-PAGE gel was fixed in fixation buffer (40% ethanol, 10% acetic acid) for 1 h, washed four times with ddH O for 20 min each, sensitized with 0.02% sodium thiosulfate for 1 min, washed three times with ddH O for 20 seconds each, incubated in ice-cold 0.1% silver nitrate solution (0.1% AgNO, 0.02% formaldehyde) for 20 min at 4 °C, washed three times with ddH O for 20 seconds each followed by one wash for 1 min, and developed with developing solution (3% sodium carbonate, 0.05% formaldehyde). The protein bands corresponding to KIN-G that were co-purified with WDR2-3HA from control and GW843682X-treated samples were excised, destained, and analyzed by mass spectrometry as described above.

### Tubulin polymerization and microtubule binding and gliding assay

*In vitro* tubulin polymerization was performed according to our published procedure (Hu et al., 2024). Non-labeled porcine brain tubulin (Cytoskeleton, Inc., Cat#: T240-A80, 80 μg) was mixed with rhodamine-labeled porcine brain tubulin (Cytoskeleton, Inc., Cat#: TL590M, 20 μg) in BRB80-DTT buffer (80 mM Potassium-PIPES, pH 6.8, 1 mM MgCl_2_, 1 mM EGTA, 1 mM DTT) supplemented with 1 mM Guanylyl-(α,β)-methylene-diphosphonate (GMP-CPP). The mixture was incubated at 4°C for 5 min and then clarified by centrifugation at 353,000 ×*g* at 4°C for 5 min in a TLA120.1 rotor in an ultracentrifuge (Beckman Coulter TL-100) to remove aggregates. The supernatant was snap-frozen in liquid nitrogen, aliquoted, and stored at −80°C. To generate microtubule seeds, an aliquot of the supernatant was diluted (1:4 v/v) with the BRB80-DTT buffer to a final concentration of 0.5 mg/ml, and incubated at 37°C for 30 min. Microtubule seeds were centrifugated at 353,000 ×*g* at 27°C for 5 min, and the pellet was resuspended in the BRB80-DTT buffer.

To polymerize microtubules from the microtubule seeds, 20 μg of rhodamine-labeled tubulin was mixed with the microtubule seeds in the presence of 1 mM GTP. Microtubules were then successively assembled by incubating at 37°C in the BRB80-DTT-GTP buffer (BRB80-DTT buffer, 1.0 mM GTP) with 0.1 μM Taxol for 20 min, 1 μM Taxol for 10 min, and 10 μM Taxol for 20 min. Assembled microtubules were diluted with 10 μM Taxol in the BRB80-DTT-GTP buffer.

Microtubule-binding and -gliding assay was conducted in a chamber assembled on a glass slide with two double-sided tapes, which were spaced 8 mm apart, and covered with a 22 mm × 30 mm cover glass. Twenty microliters of 3 μM recombinant KIN-G-6His or its mutants, which were suspended in the BRB80-DTT-ATP-Taxol buffer containing 800 ng/ml BSA, was loaded into the chamber and incubated for 3 min to allow the attachment of KIN-G-6His to the cover glass. The chamber was washed twice with 20 μl of the BRB80-DTT-ATP-Taxol buffer, and 20 μl of 120 nM rhodamine-labeled microtubules in the BRB80-DTT-ATP-Taxol buffer containing 2.25 mg/ml BSA was then loaded into the chamber. After incubation for 2 min, the chamber was observed for microtubule binding and gliding under the Nikon A1 confocal microscope, and images were captured every 5 seconds. Movie files of the AVI format were generated by NIS Elements (Nikon).

To test the effect of TbPLK phosphorylation on KIN-G microtubule binding and gliding, two experiments were performed. In experiment #1, 4 μM recombinant KIN-G-6His was pre-incubated with 2 μM GST-TbPLK-6His or GST-TbPLK^K70R^-6His in the presence of 1 mM ATP at 37°C for 7 min, and the protein mixture was clarified by centrifugation at 21,000 ×*g* for 30 sec. The supernatant was used for microtubule-binding and - gliding assay as described above. In experiment #2, 4 μM recombinant KIN-G-6His was settled onto glass coverslip in the chamber. Microtubules were then added into the chamber, and incubated for 5 min. Subsequently, 0.2 μM GST-TbPLK-6xHis or the BRB80-DTT-ATP-Taxol buffer was added into the chamber, and image acquisition was started after 2 min of incubation in the Nikon A1 confocal microscope.

Microtubules bound to KIN-G or its mutants were quantified in the last image of the time-lapse imaging process or at the indicated time point of the time-lapse imaging process. Nonspecifically coverslip-attached microtubules, which showed no gliding during the time-lapse imaging process, were not counted as KIN-G-bound microtubules and, thus, were excluded. Microtubule-gliding speed was calculated using the microtubules that exhibited continuous movement for at least 100 sec (20 frames) in time-lapse imaging.

## Acknowledgements

We are grateful to Dr. Cynthia Y. He of National University of Singapore for providing anti-CC2D antibody and anti-TbGRASP antibody, Dr. Derrick Robinson of University of Bordeaux for providing anti-TbBILBO1 antibody, and Dr. Hira Nakhasi of the FDA for providing anti-TbCentrin4/LdCen1 antibody. This work was supported by the NIH R01 grants AI118736 and AI101437 to Z. L. The funders do not play any role in the study design, data collection and analysis, decision to publish, or preparation of the manuscript. The image of microtubules in Figure 8 was created in BioRender (https://BioRender.com/b6kkzaq).

## Competing interests

The authors declare that they have no conflicts of interest with the contents of this article.

---

**Figure 2-video 1: Microtubule-binding and -gliding activities of KIN-G.**

**Figure 2-video 2: Microtubule-binding and -gliding activities of KIN-G, which was pre-incubated with TbPLK.**

**Figure 2-video 3: Microtubule-binding and -gliding activities of KIN-G, which was pre-incubated with the kinase-dead mutant TbPLK^K70R^.**

**Figure 2-video 4: Microtubule-binding and -gliding activities of KIN-G, with TbPLK added to the chamber after microtubules were added.**

**Figure 3-video 1: Microtubule-binding and -gliding activities of KIN-G.**

**Figure 3-video 2: Microtubule-binding and -gliding activities of KIN-G^T301D^.**

**Figure 3-video 3: Microtubule-binding and -gliding activities of KIN-G^T301A^.**

**Figure 3-video 4: Microtubule-binding and -gliding activities of KIN-G^T284D^.**

**Figure 3-video 5: Microtubule-binding and -gliding activities of KIN-G^T284A^.**

**Figure 1-source data 1. PDF file containing original western blots, Coomassie blue-stained gel image and silver-stained gel image for Figure 1B, 1C, and 1D, indicating the relevant bands and treatments.**

**Figure 1-source data 2. Original files for western blot analysis, Coomassie blue staining, and silver staining displayed in Figure 1B, 1C, and 1D.**

**Figure 4-source data 1. PDF file containing original western blots for Figure 4B, indicating the relevant bands and treatments.**

**Figure 4-source data 2. Original files for western blot analysis displayed in Figure 4B**.

## Notes

### Competing Interest Statement

The authors have declared no competing interest.

### Summary of Updates

Figure 8 revised. Supplemental files updated.

