## Supplementary material for "Polo-like kinase phosphorylation of the orphan kinesin KIN-G negatively regulates centrin arm biogenesis in *Trypanosoma brucei*": Figure 1-source data-1

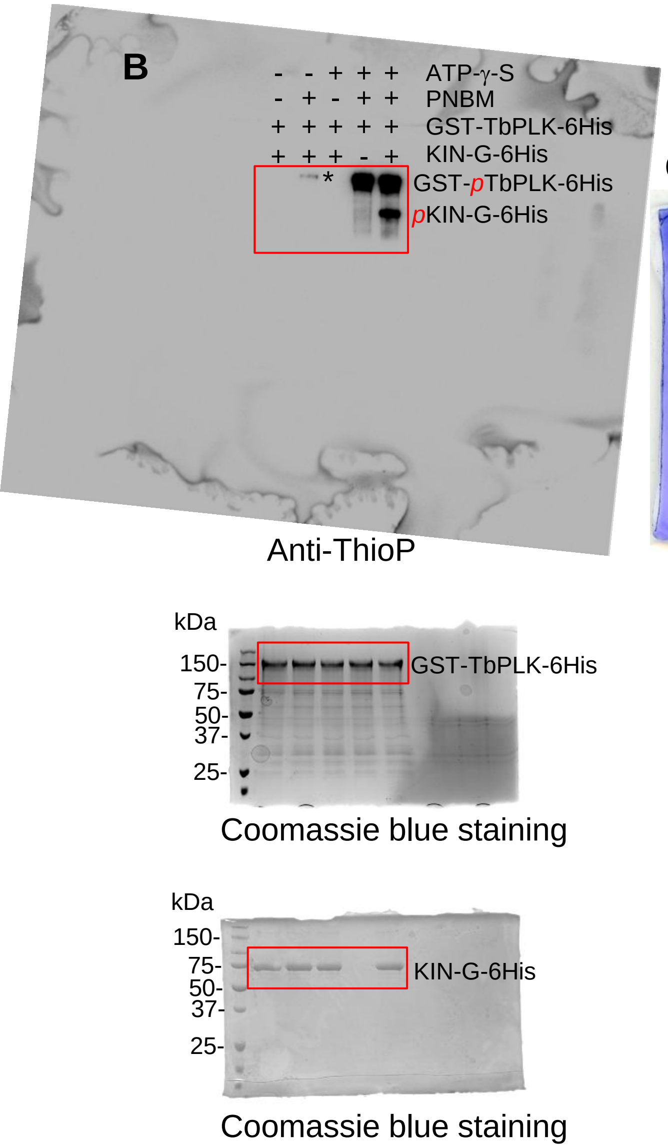


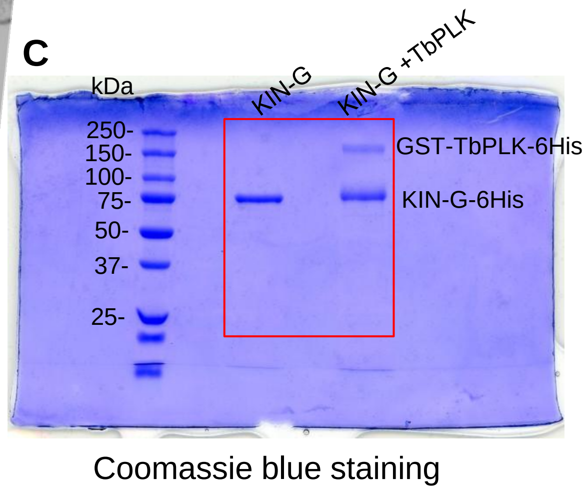


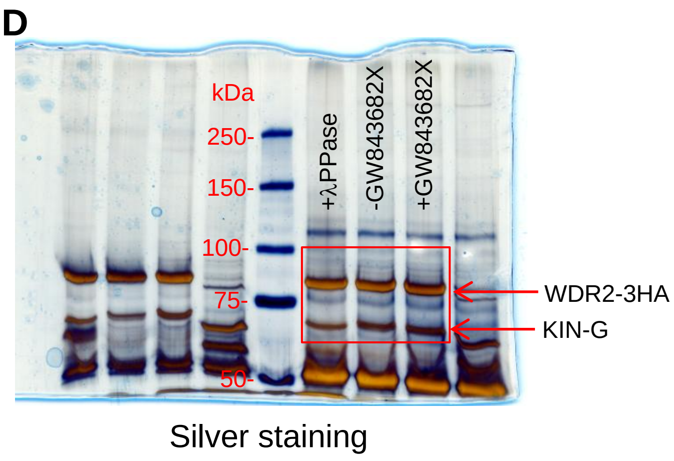


**Figure 1 - source data 1.** Original western blots, Coomassie-stained gels and silver-stained gels, corresponding to Figure 1, panel B, panel C, and panel D.
