## Supplementary figures and images for "Polo-like kinase phosphorylation of the orphan kinesin KIN-G negatively regulates centrin arm biogenesis in *Trypanosoma brucei*"

### Figure 4-source data-1

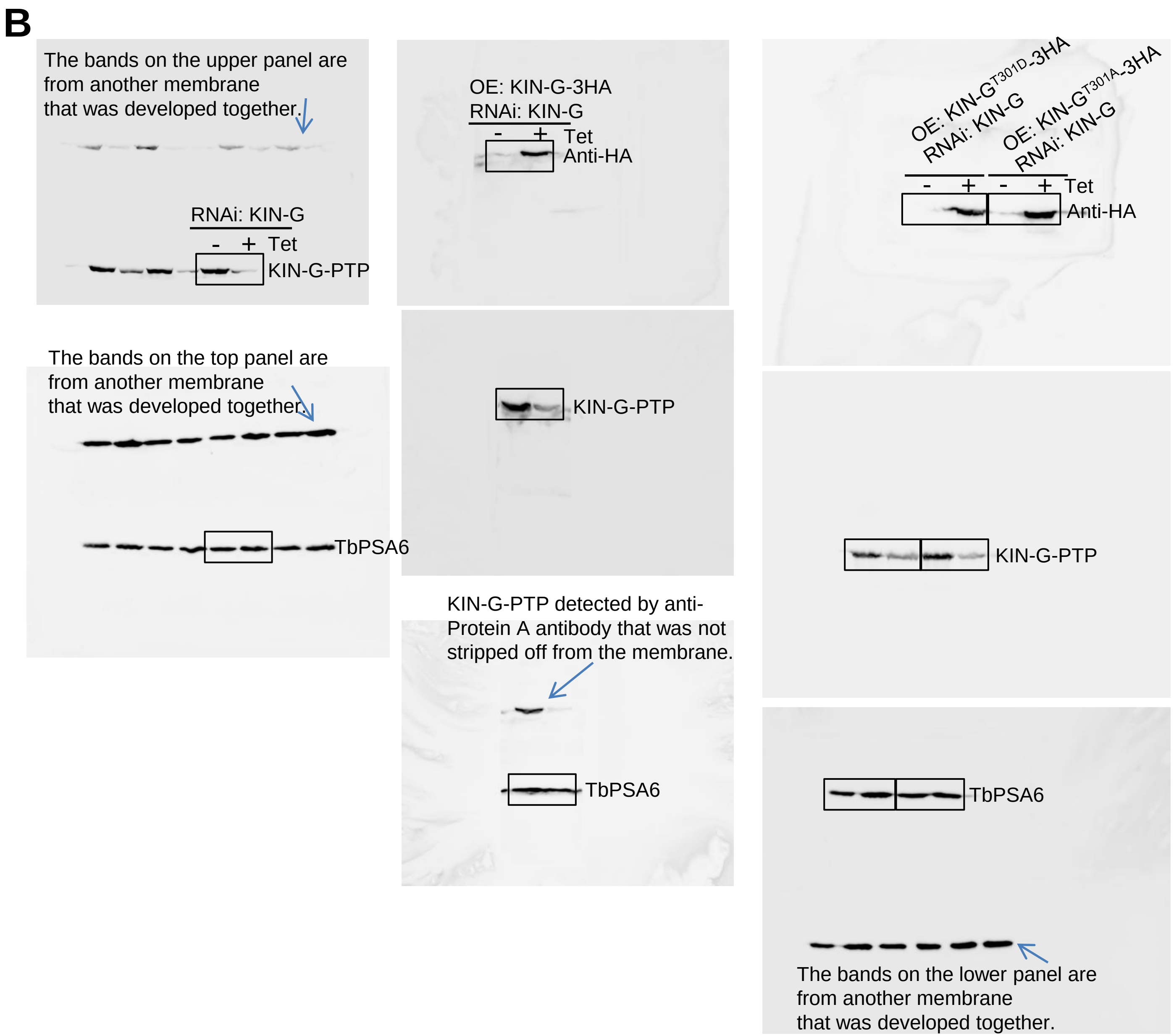


**Figure 4 - source data 1.** Original western blots corresponding to Figure 4, panel B.

### GST-TbPLK.tif

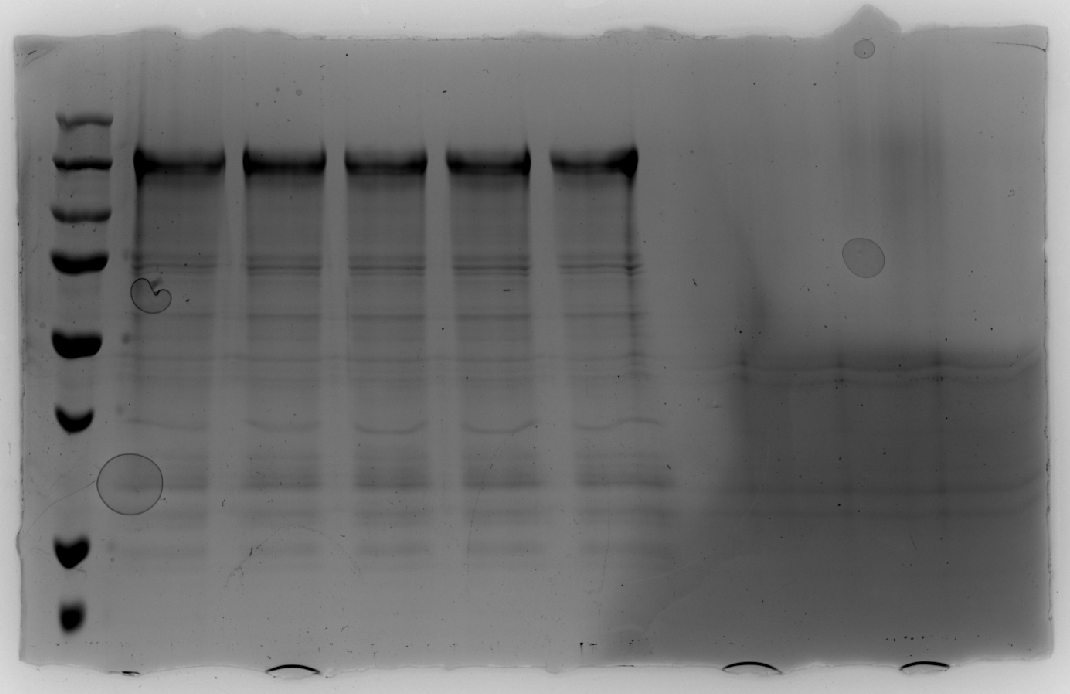

### KIN-G RNAi_KIN-G-PTP_anti-Protein A immunoblotting.tif

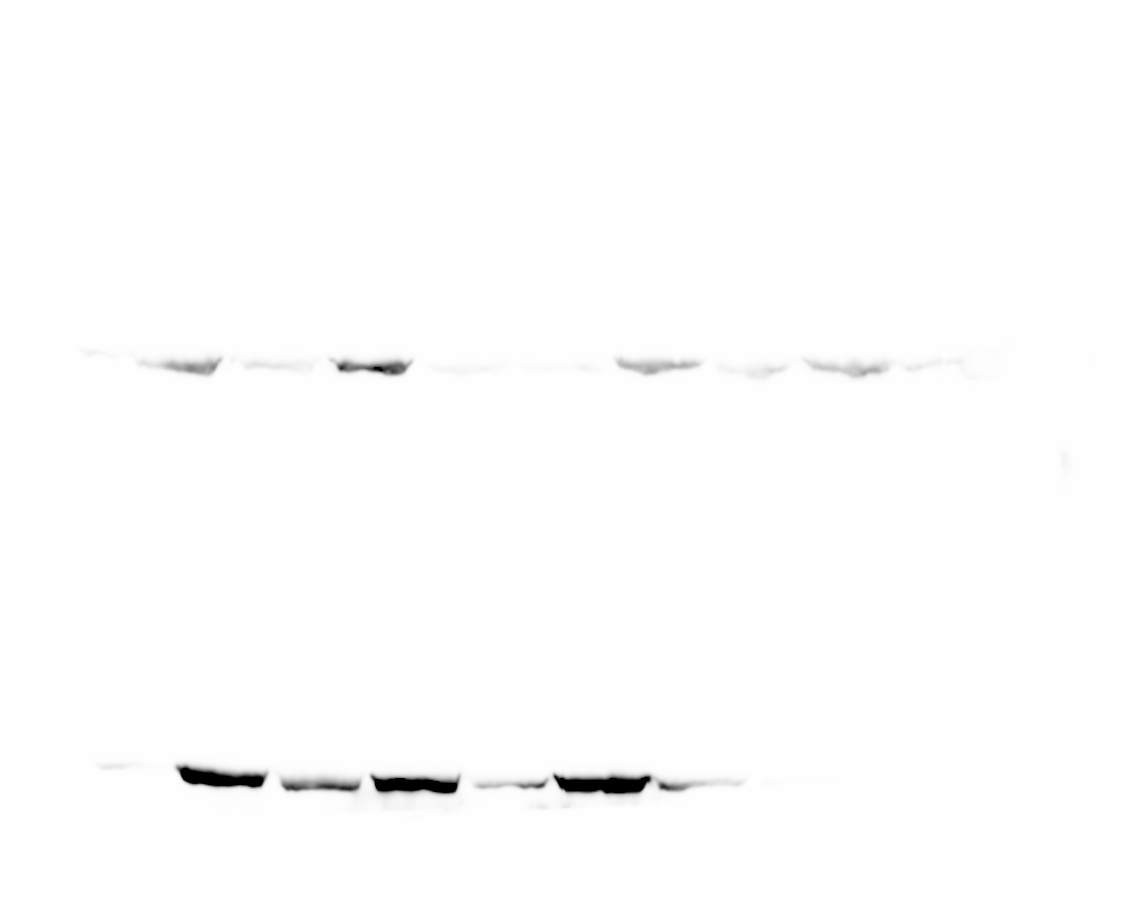

### KIN-G RNAi_KIN-G-PTP_anti-TbPSA6 immunoblotting.tif

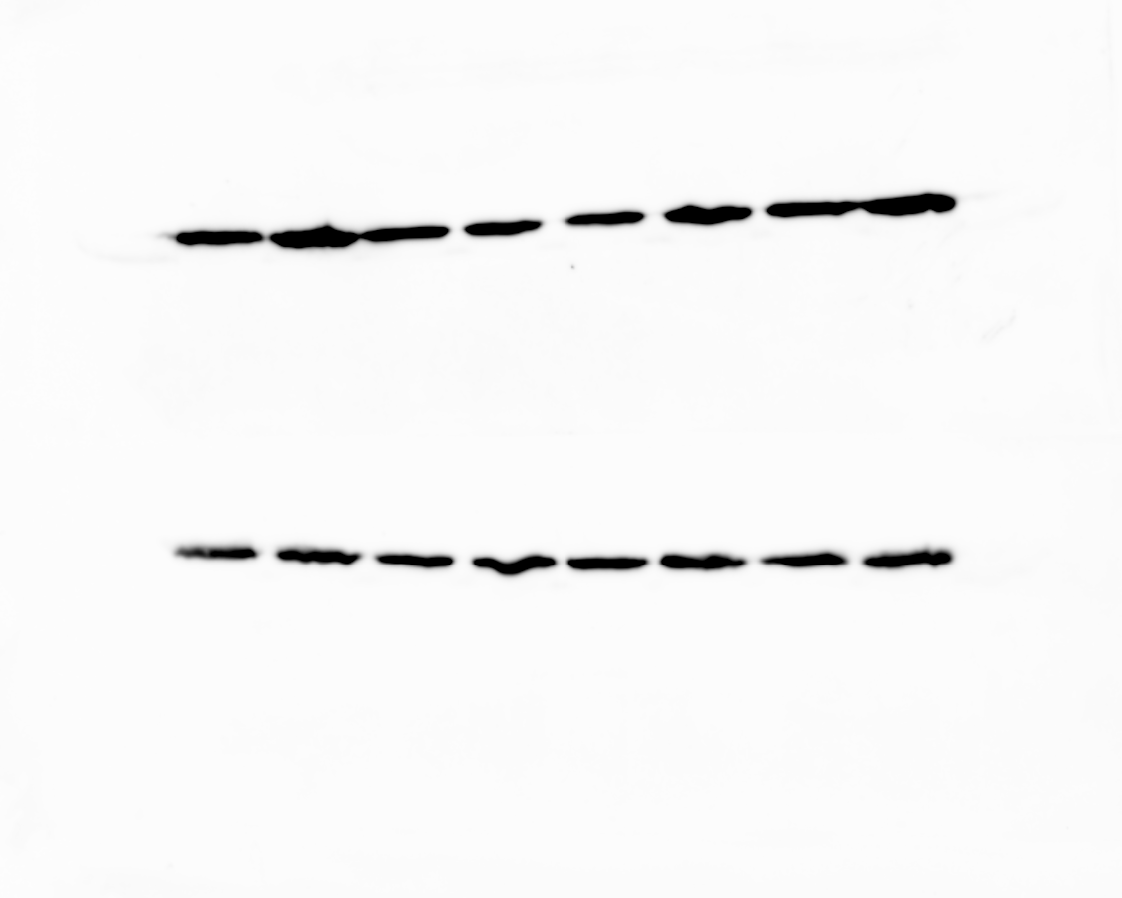

### KIN-G RNAi_KIN-G-PTP_KIN-G OE_anti-Protein A immunoblotting.tif

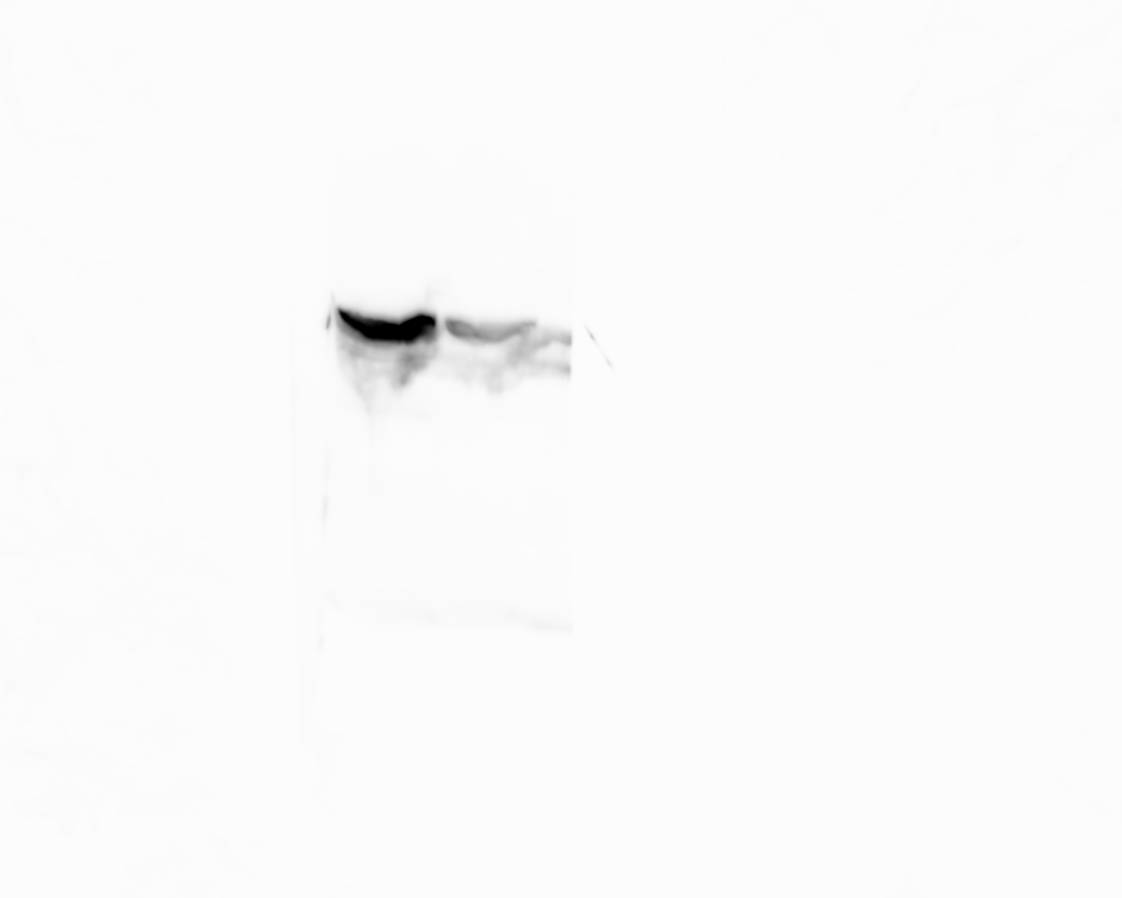

### KIN-G RNAi_KIN-G-PTP_KIN-G-3HA OE_anti-HA immunoblotting.tif

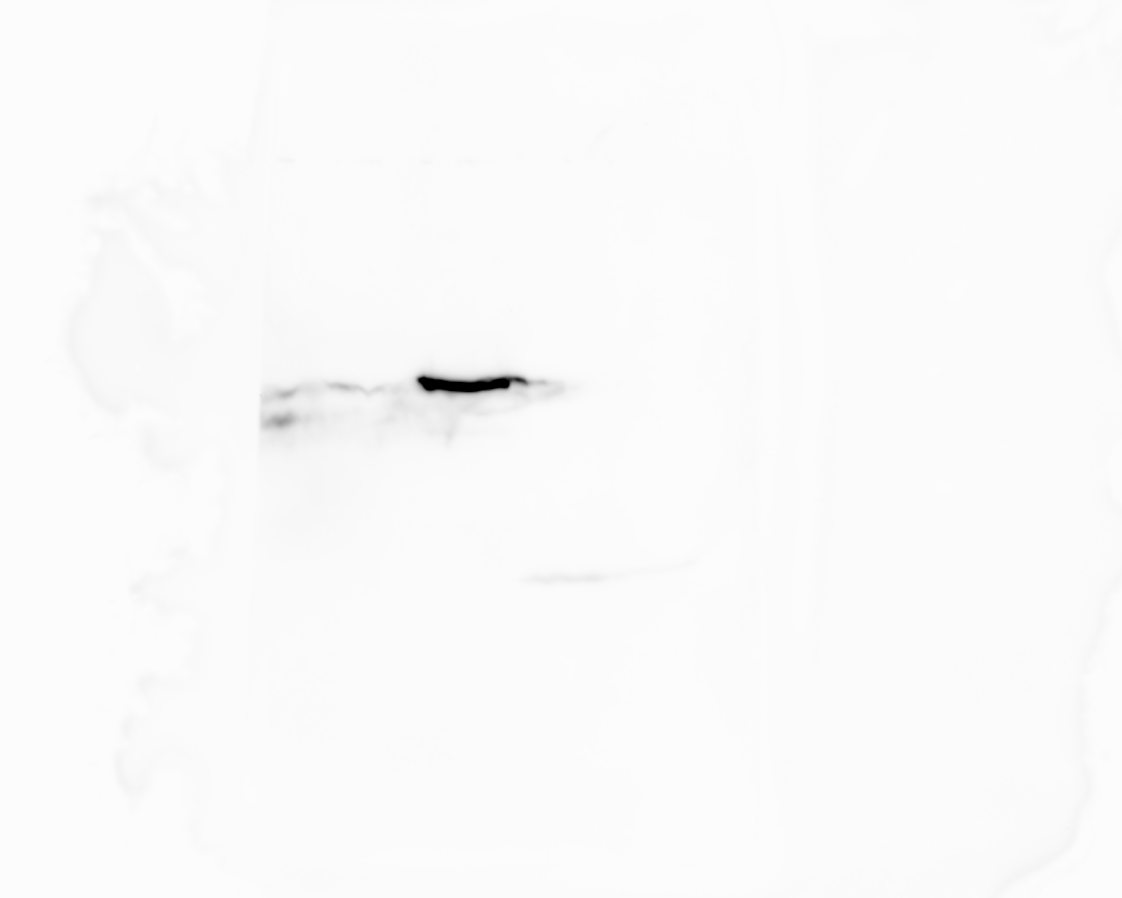

### KIN-G RNAi_KIN-G-PTP_KIN-G-3HA OE_anti-TbPSA6 immunoblotting.tif

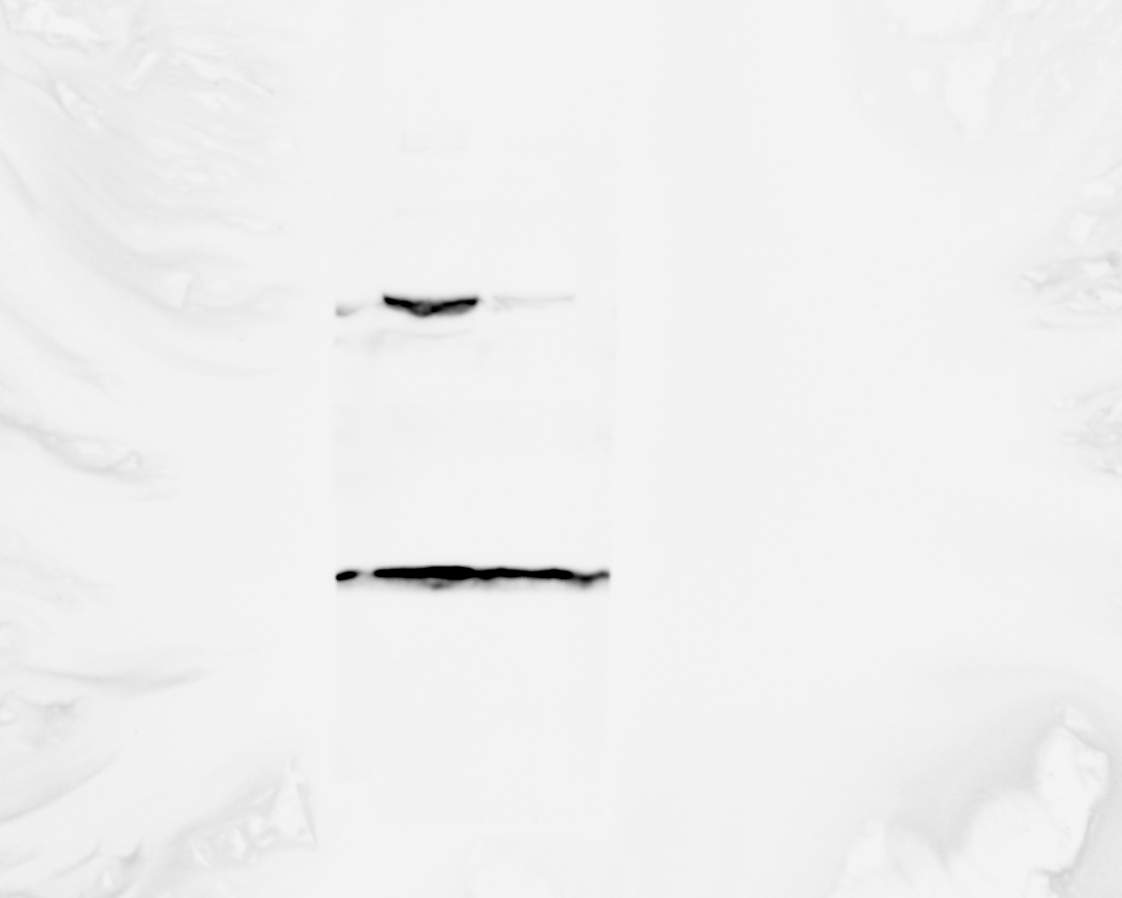

### KIN-G RNAi_KIN-G-PTP_KIN-G-T301D-3HA or T301A-3HA OE_anti-HA immunoblotting.tif

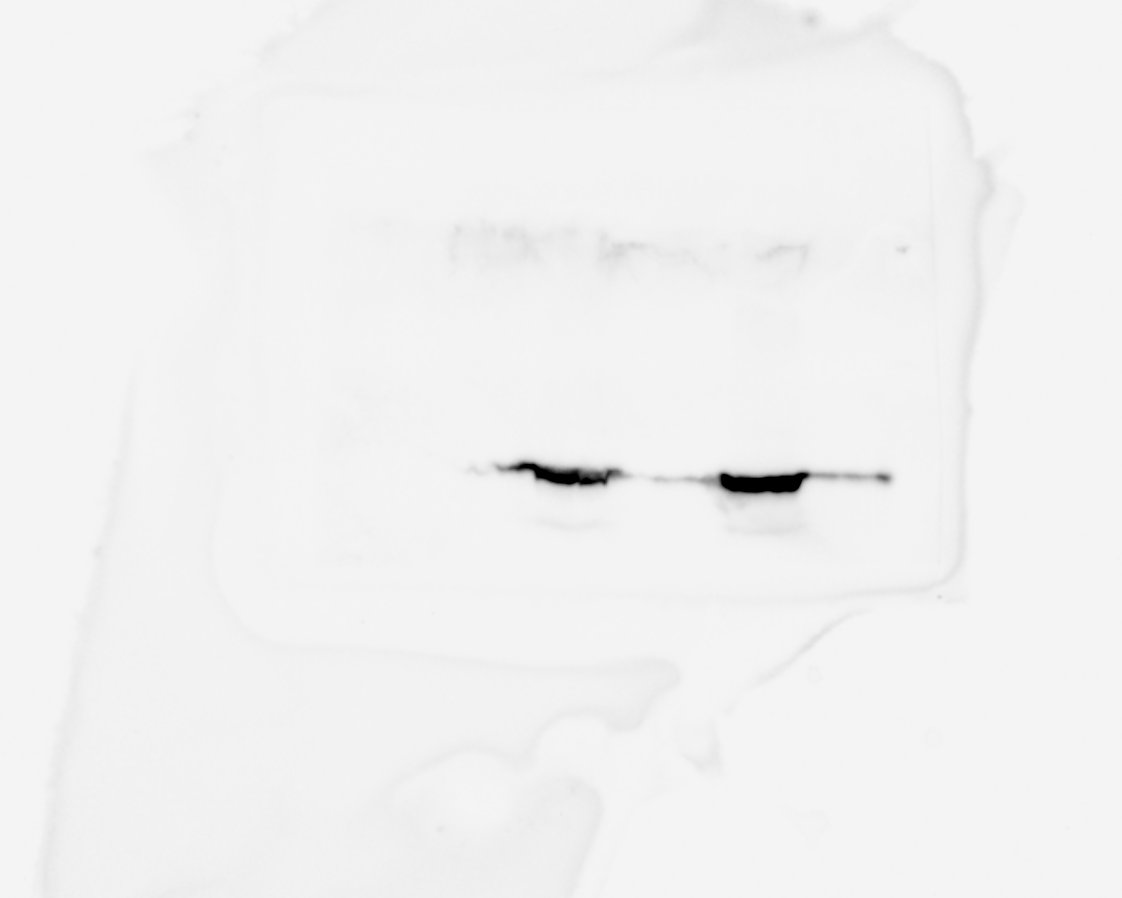

### KIN-G RNAi_KKIN-G-PTP_KIN-G-T301D-3HA or T301A-3HA OE_anti-Protein A immunoblotting.tif

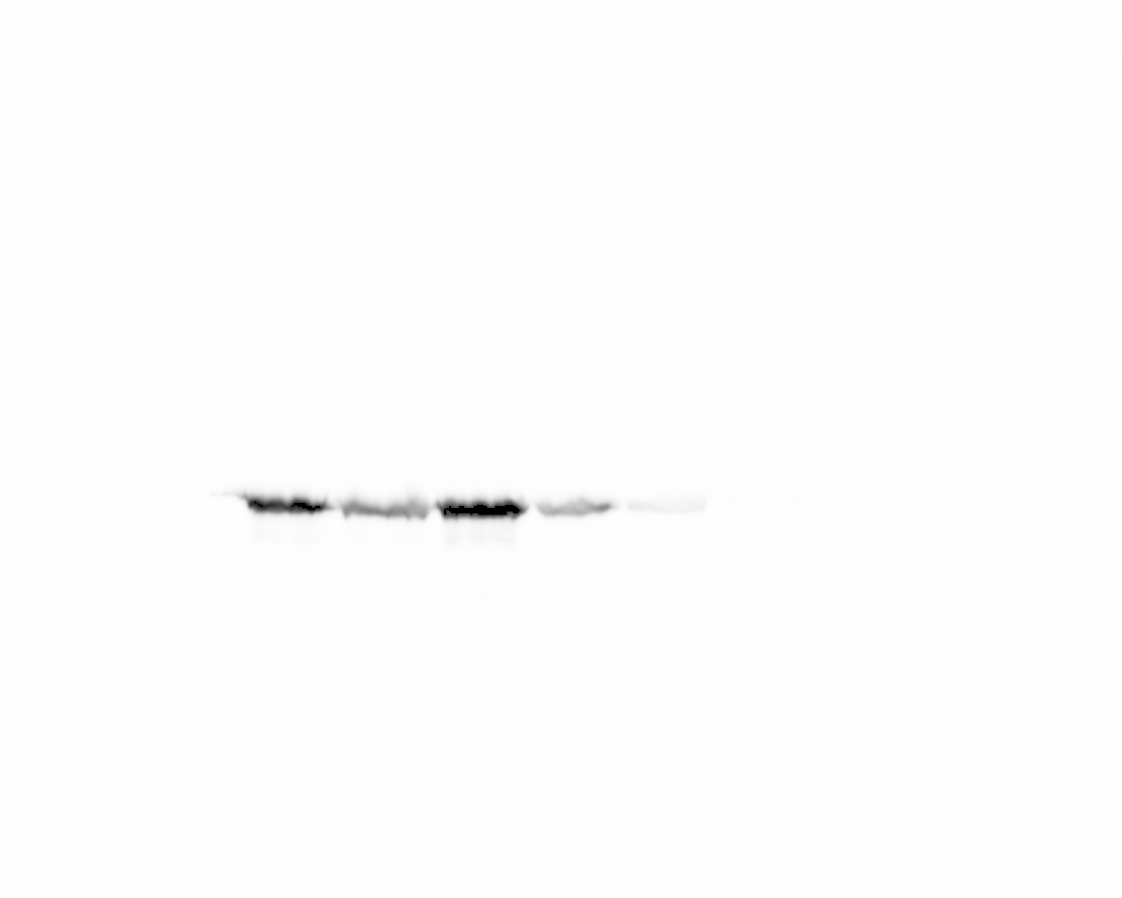

### KIN-G RNAi_KKIN-G-PTP_KIN-G-T301D-3HA or T301A-3HA OE_anti-TbPSA6 immunoblotting.tif

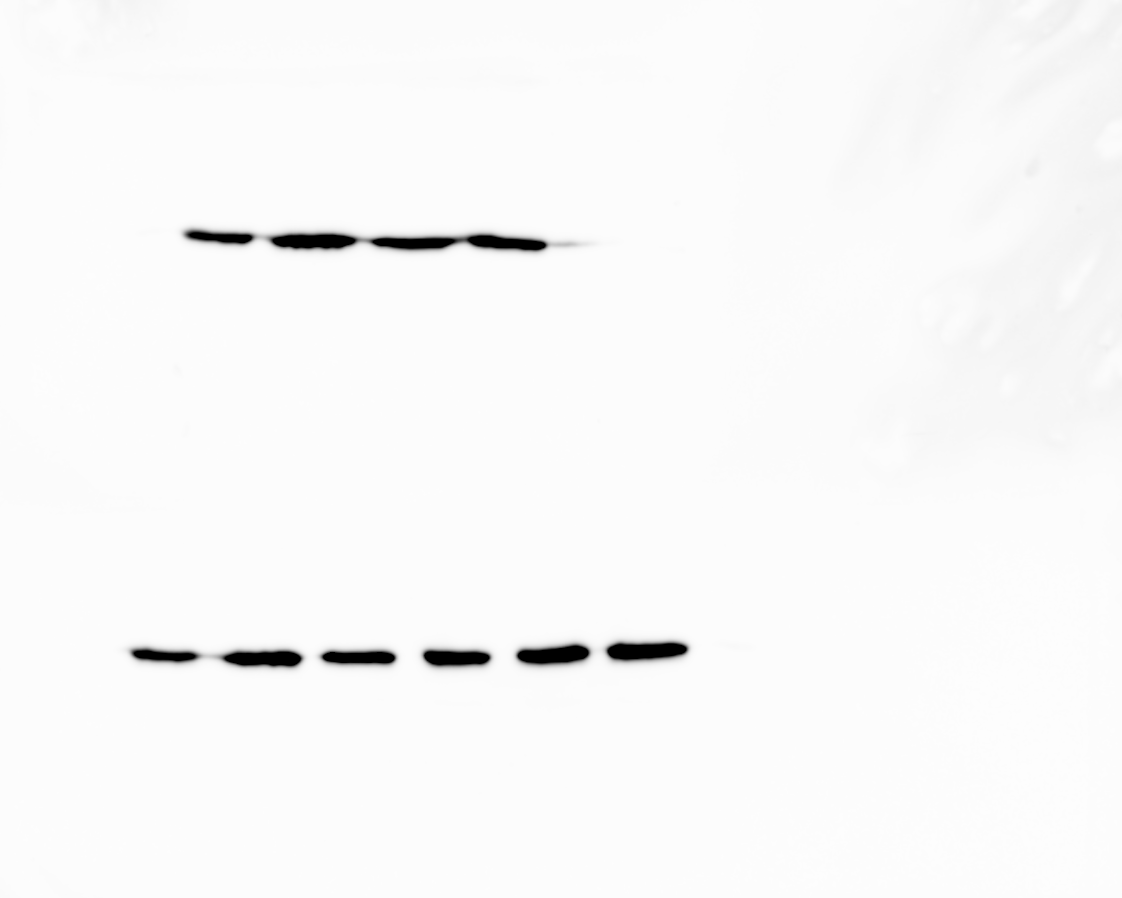

### KIN-G-6xHis.tif

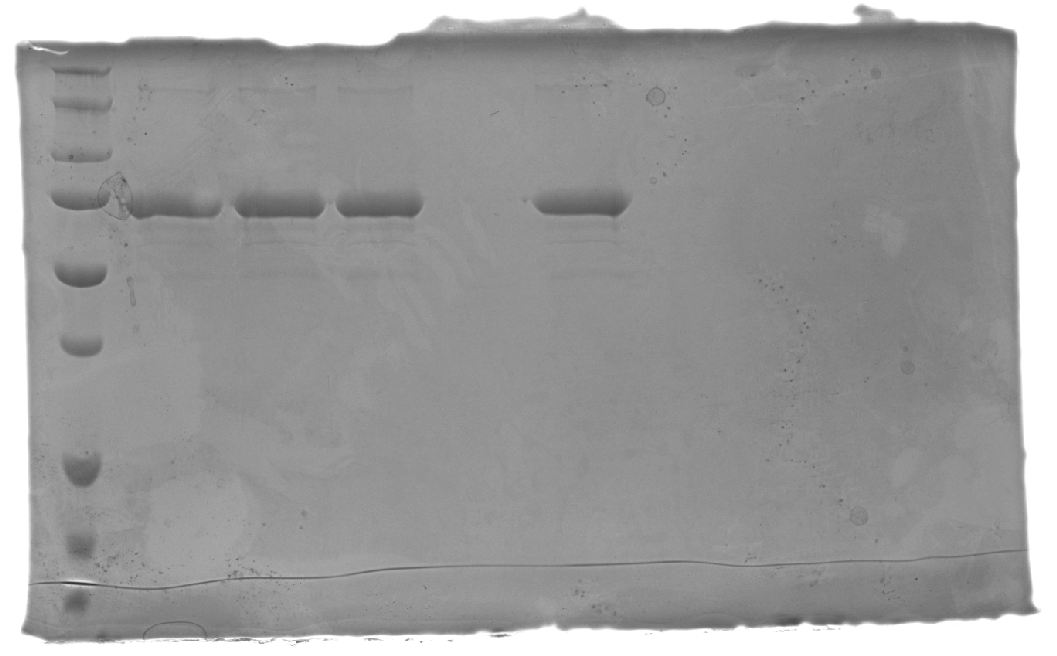

### KIN-G-TbPLK kinase assay_ anti-ThioP.tif

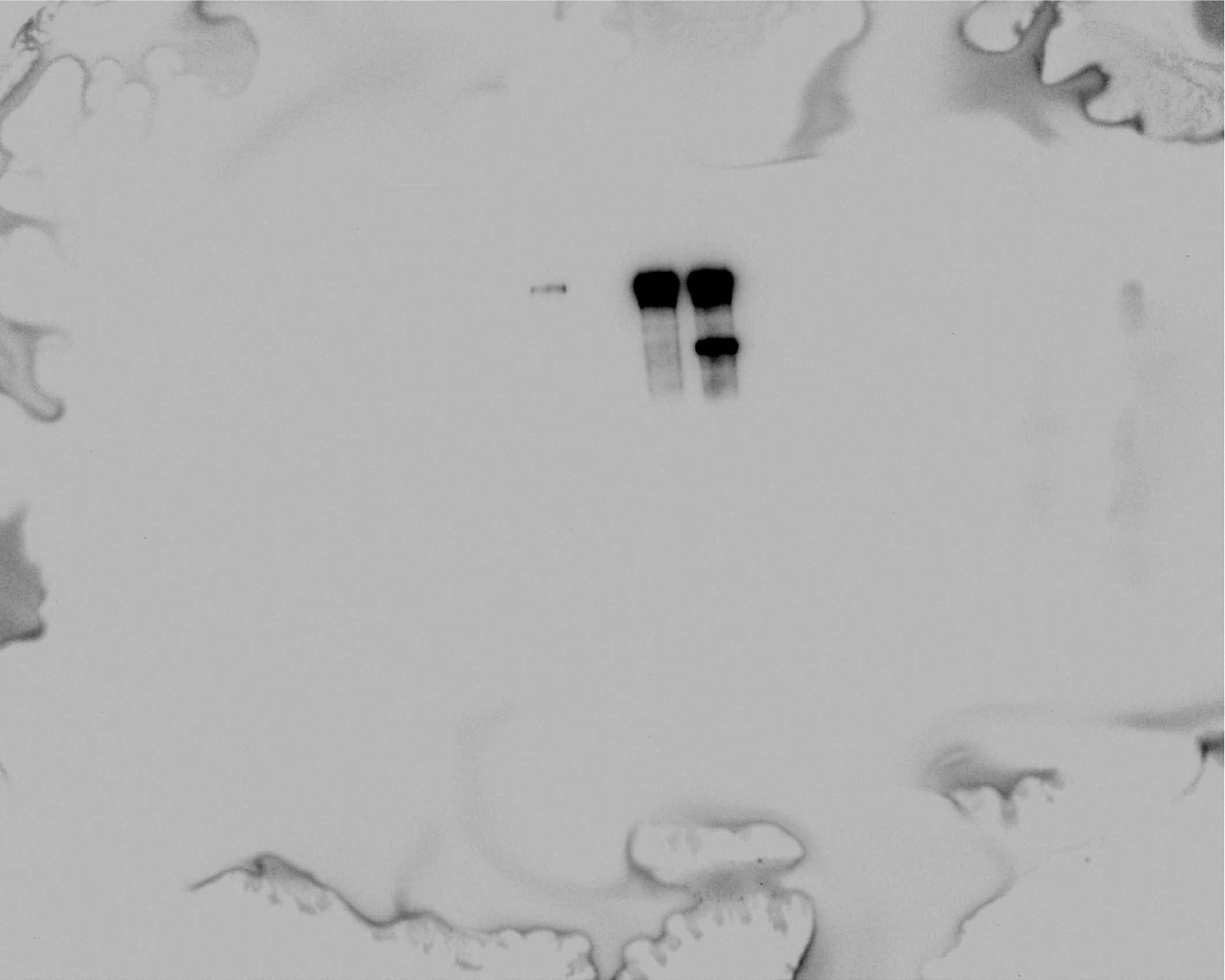

### KIn-G-TbPLK kinase assay_Commassie blue staining.tif

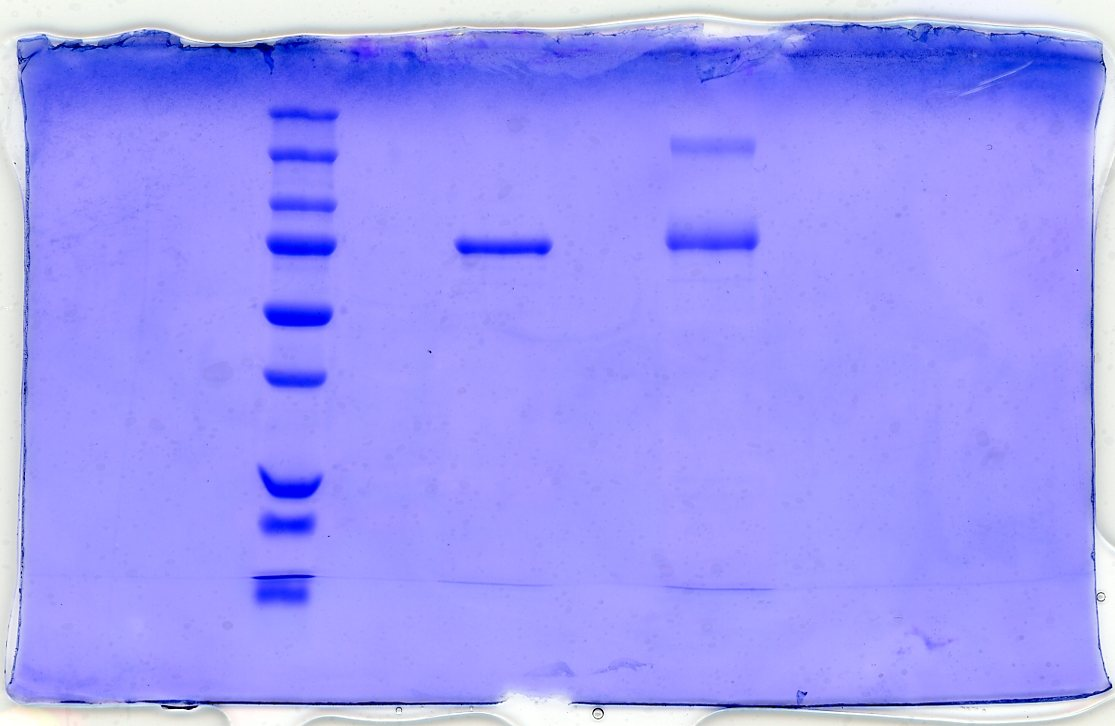

### WDR2-3HA IP_silver staining.tif

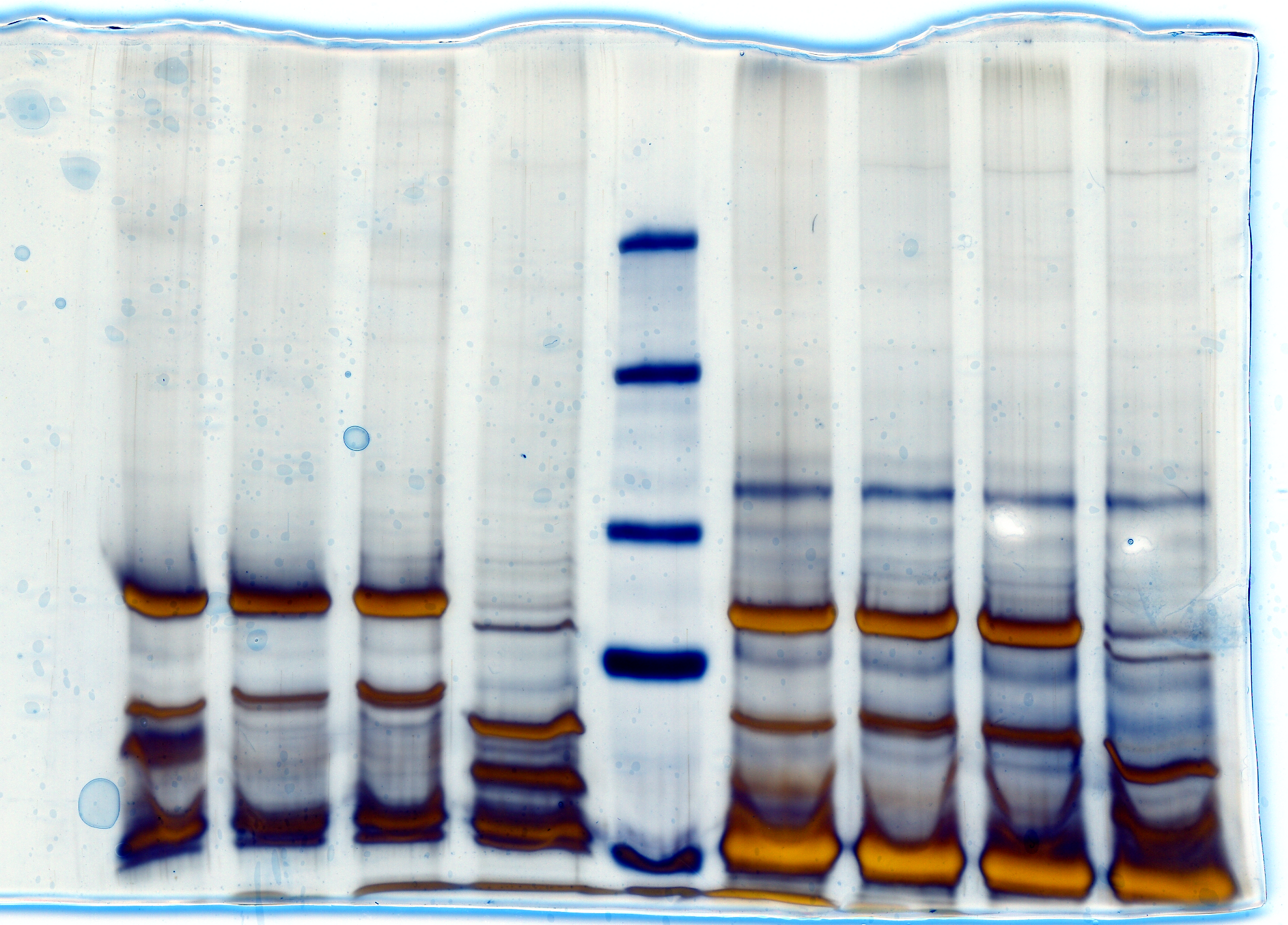
